# Molecular basis for substrate recruitment to the PRMT5 methylosome

**DOI:** 10.1101/2020.08.22.256347

**Authors:** Kathleen M. Mulvaney, Christa Blomquist, Nischal Acharya, Ruitong Li, Meghan O’Keefe, Matthew Ranaghan, Matthew Stokes, Alissa J Nelson, Sidharth S. Jain, Josie Columbus, Fazli K. Bozal, Adam Skepner, Donald Raymond, David C. McKinney, Yelena Freyzon, Yossef Baidi, Dale Porter, Alessandra Ianari, Brian McMillan, William R. Sellers

## Abstract

PRMT5 is an arginine methyltransferase and a therapeutic target in *MTAP* null cancers. PRMT5 utilizes adaptor proteins for substrate recruitment through a previously undefined mechanism. Here, we identify an evolutionarily conserved peptide sequence shared among the three known substrate adaptors (pICln/CLNS1A, RIOK1 and COPR5) and show it is necessary and sufficient for interaction with PRMT5. We structurally resolve the interface with PRMT5 and show via genetic perturbation that it is required for methylation of adaptor-recruited substrates including the spliceosome, histones, and ribosome assembly complexes. Genetic disruption of the PRMT5-substrate adaptor interface leads to a hypomorphic decrease in growth of *MTAP* null tumor cells and is thus a novel site for development of therapeutic inhibitors of PRMT5.

---

Protein arginine methyltransferase 5 (PRMT5) is a Type II arginine methyltransferase and an essential enzyme^1–3^. PRMT5 symmetrically dimethylates substrate proteins by catalyzing the 2-step transfer of 2 methyl groups from 2 S-adenosyl methionine (SAM) co-factor molecules to specific arginine residues within a substrate^4,5^. Through substrate methylation, PRMT5 regulates chromatin structure, gene transcription, cellular differentiation, and mRNA splicing^2,5–8^. Substrate methylation depends upon assembly of a larger PRMT5 methylosome complex that includes its obligate binding partner WDR77. Together, PRMT5 and WDR77 form a stable 4:4 heterooctamer complex^9,10^. The PRMT5 methylosome requires additional partners (referred to as substrate adaptors herein) to recognize and methylate substrates.

There are three known PRMT5 substrate adaptors: pICln/CLNS1A, RIOK1, and COPR5. pICln, a chaperone protein that regulates assembly of the splicesome, recruits PRMT5 to subunits of the splicesome for methylation, including SmB/B’, SmD1, and SmD2^11–13^, as well as ribosomal proteins LSM4, LSM10 and LSM11^14–16^. Through these interactions, PRMT5 regulates cellular splicing activity^17^ and the formation of RNA processing bodies^15^. RIOK1, an atypical serine/threonine kinase, recruits nucleolin and RPS10 to the PRMT5 complex for methylation^18^. Through this complex PRMT5 regulates ribosome biogenesis^19^. Lastly, COPR5 recruits PRMT5 to nucleosomes and promotes methylation of histone arginine residues, including H3R8 and H4R3^20^. These methyl marks regulate recruitment of chromatin remodeling complexes^7,20,21^. While RIOK1 and pICln compete for PRMT5 binding^18^, the precise mechanism by which substrate adaptors associate with PRMT5 has not been resolved.

We and others identified PRMT5 as a cancer dependency in *MTAP* deleted tumors^22–24^. Specifically, shRNA-induced depletion of PRMT5 or either of the substrate adaptors RIOK1 and pICln confer synthetic lethality in this setting. The *MTAP* gene is adjacent to the frequently deleted tumor suppressor locus containing CDKN2A/B (p16/p14) and is thus co-deleted in ~15% of all human tumors. In *MTAP* null tumors, deficiency of MTAP (methylthioadenosine phosphorylase) results in accumulation of its substrate, methylthioadenosine (MTA). Increased levels of MTA result in partial inhibition of the PRMT5 enzyme by competing with SAM for the co-factor binding site^22^. PRMT1 and MAT2A (involved in SAM biogenesis) are also top correlates of PRMT5 in the cancer dependency map in MTAP null cancers^1^. Finally, leukemia and lymphoma models are sensitive to PRMT5 inhibitors independent of MTAP status^25–28^. Based on these data, direct PRMT5 catalytic inhibitors as well as PRMT5 pathway-related inhibitors (i.e. MAT2A or PRMT1 inhibitors) are being tested in a total of five human clinical trials in both the MTAP-deficient and MTAP-proficient settings^29^.

Here, we identify a novel mechanism by which PRMT5 interacts with its adaptor proteins. Each substrate adaptor protein contains a highly conserved linear peptide sequence, here termed the PRMT5 binding motif (PBM), which is both necessary and sufficient for PRMT5 interaction. The PBM binding surface on PRMT5 is distal to the catalytic site, but is required for methylation of many well characterized substrates. Mutations in this PRMT5 site or in the complementary PBM motif impair viability in *MTAP* null cancers. Thus, we have identified a novel mechanism through which PRMT5 recruits substrates and a potentially druggable functional site on the methylosome.

## Results

### Identification of a PRMT5 Binding Motif (PBM)

PRMT5 recruits many well characterized substrates through substrate adaptor proteins. To elucidate the mechanism(s) for the PRMT5:adaptor protein interaction, the primary amino acid sequences and secondary/tertiary structures between the three adaptor proteins were assessed. Amino acid sequence alignment of the three PRMT5 adaptor proteins pICln, COPR5 and RIOK1 revealed a linear peptide sequence common to each substrate adaptor protein: GQF[D/E]DA[D/E] (Figure 1A). For pICln and RIOK1, this peptide sequence is conserved across chordates (Figure 1A). COPR5 protein homologs are present only in tetrapods with the peptide sequence conserved across all classes. These sequences (Figure 1A, red block) reside near the N or C-terminus of these proteins and lie outside of recognized Pfam domains.

**Figure 1.**
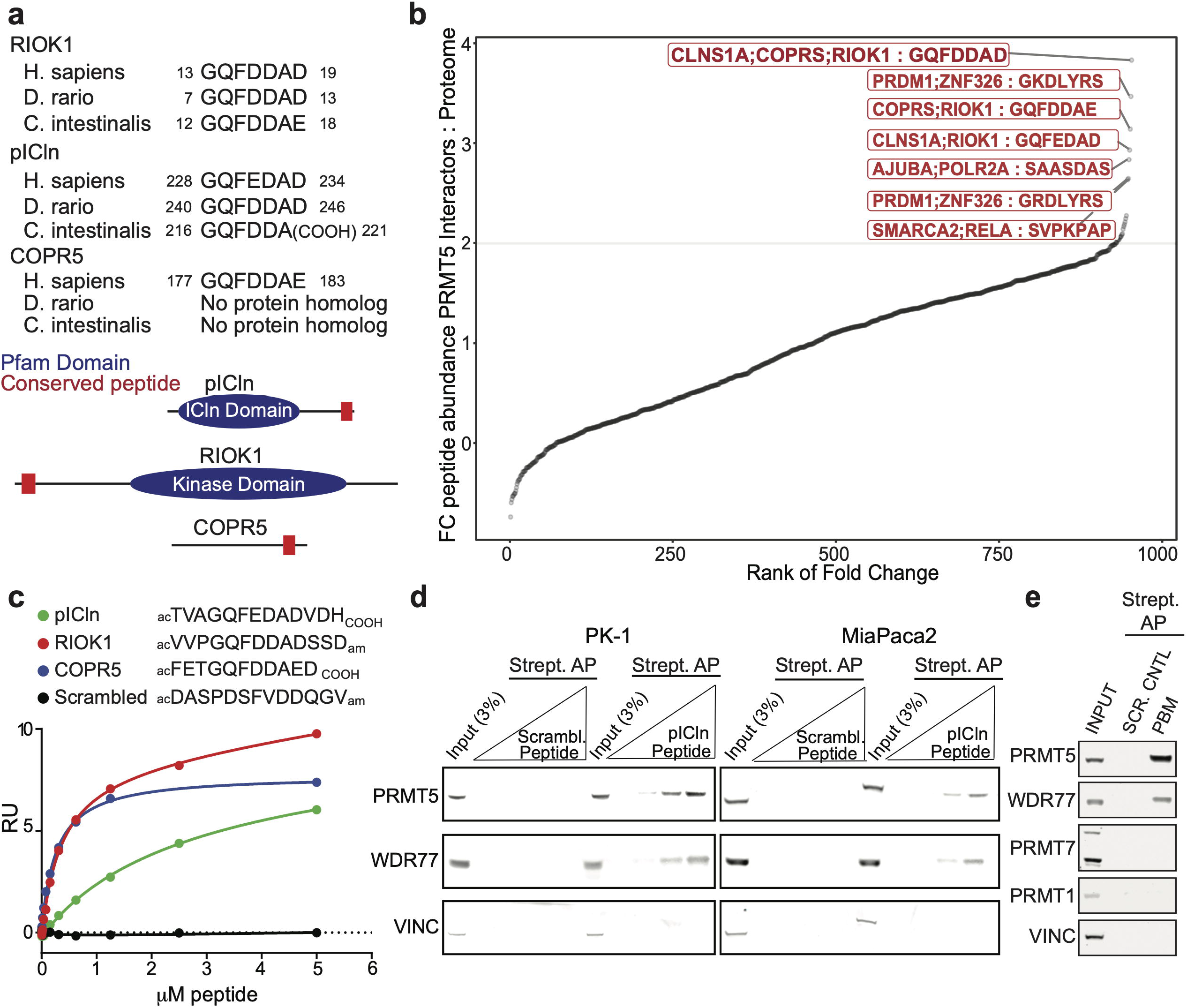
Identification of a conserved PRMT5 substrate adaptor binding motif: GQF[D/E]DA[D/E]. a. Amino acid sequence alignment of three known substrate adaptor proteins of PRMT5 (COPR5, pICln, and RIOK1) identified an evolutionarily conserved 7-mer peptide that lies at the N-terminus of RIOK1 and C-termini of pICln and COPR5. b. Peptide distribution of 7 amino acid sequence motifs across all PRMT5 binding proteins (BioGrid) is shown. GQF[D/E]DA[D/E] was enriched in PRMT5 binding proteins compared to other peptide sequences comparing 7 amino acid length peptides in PRMT5 interactors to 7-mer peptides across the human proteome. The BLOSUM Clustered Scoring Matrix was utilized to maximize the flexibility of a potential functional protein pocket, i.e. allowing for D/E at the same position. c. Interactions between the PRMT5 complex and the conserved 7-mer peptides described in (A) were measured by SPR. The PRMT5 complex was immobilized by biotin-neutravidin and the indicated peptides titrated as analyte. The RU response at equilibrium is shown. d. Streptavidin affinity purification (AP) from cell lysates derived from MiaPaca2 and PK1 cells that were incubated with increasing concentrations of either the pICLn 17-mer peptide containing the 7-mer sequence in (A) or a scrambled control peptide. Immunoblots show the amount of PRMT5 and WDR77 that were co-purified with the peptides. Vinculin serves as a loading and affinity purification control. e. Streptavidin AP from MiaPaca2 cell lysates that were incubated with 1μg of either the pICLn 17-mer peptide (“PBM”) containing the 7-mer sequence in (A) or a scrambled control peptide. Immunoblots show the amount of PRMT5, WDR77, PRMT1 and PRMT7 that were co-purified with the peptides. Vinculin serves as a loading and AP control.

To determine whether this 7-mer peptide is unique to PRMT5 substrate adaptors, the frequency of all 7-amino acid sequences was counted among PRMT5 interactors determined in BioGrid and among non-interactors. Using a BLOSUM Clustered Scoring Matrix to maximize the flexibility of a potential functional protein pocket (i.e. allowing for D/E at the same position)^30^, GQFDDAD was found as the most enriched peptide among PRMT5 interacting proteins (Figure 1B) and the GQF[D/E]DA[D/E] peptide sequence is unique to pICln, COPR5, and RIOK1 across the proteome (Figure 1B). Given the conservation of the peptide among adaptor proteins and its uniqueness within the proteome, we hypothesized that this site might mediate substrate adaptor interaction with PRMT5.

### The PRMT5 Binding Motif is sufficient to bind the PRMT5 methylosome

To determine whether the adaptor peptide sequence is sufficient to bind PRMT5, surface plasmon resonance (SPR) was performed using purified PRMT5:WDR77 complex and synthetic peptides including the 7-mer sequence and a small flanking region from each adaptor (Figure 1C). Binding was detected for each of the 3 peptides, whereas no interaction was detected for a scrambled control peptide (Fig. 1C). The RIOK1 and COPR5 peptides have similar affinity for the immobilized methylosome (K_D_ = 0.35 and 0.27 μM, respectively) while the interaction between the pICln peptide and PRMT5 is weaker (K_D_ = 2.5 μM). These data demonstrate that the 7-mer peptide is sufficient for in vitro binding to PRMT5.

To determine whether the adaptor peptide is sufficient to bind endogenous PRMT5, biotinylated pICln or control peptides were incubated with extracts from two human cell lines (MiaPaca-2 and PK-1). Biotinylated peptides and bound proteins were affinity purified and PRMT5 and WDR77 detected by immunoblot. In contrast to control peptides, pICln peptides co-purified PRMT5 and WDR77 in a concentration-dependent manner (Figure 1D). In parallel experiments, co-purification of PRMT5 and WDR77, but not PRMT7 or PRMT1 was observed (Figure 1E). These data suggest that the pICln peptide can bind specifically to the endogenous PRMT5 methylosome complex from cells (Figure 1D). Henceforth, we refer to this amino acid sequence as the PRMT5-binding motif or PBM and the respective peptides as PBM peptides.

### The PBM associates with an exposed groove within the TIM barrel of PRMT5

Next, crystals of the PRMT5:WDR77 complex were generated as previously described^31^ and soaked with PBM peptides from RIOK1 (_ac_VVPGQFDDADSSD_am_) or pICln (_ac_TVAGQFEDADVDH_COOH_) and the resulting structures were resolved to 2.11 and 2.86 Å, respectively (PDB IDs 6V0N and 6V0O, respectively, Supplemental Table 1). As previously observed, the asymmetric crystal unit contains one protomer unit (a single heterodimer of PRMT5:WDR77)^10^. The full hetero-octamer (4x protomer) complex, as observed in solution, is reconstructed via crystal symmetry. The peptide-bound structures were alternatively solved at lower symmetry, but no differences between the individual protomers were observed.

Unambiguous electron densities for the peptides were observed at the TIM barrel domain of PRMT5 (Figures 2A and S1). The overall pose of the RIOK1 and pICln peptides are identical (Figure 2B) with an overall RMSD of 0.3 Å. Outside of the PBM binding site, the PRMT5 and WDR77 structures are unchanged from previous observations^9,22^. The PBM binding site is conserved across PRMT5 homologs in comparison to the protein as a whole (Figure 2A), albeit to a lesser degree than the nearly universally conserved catalytic pocket residues of the Rossmann fold.

**Figure 2.**
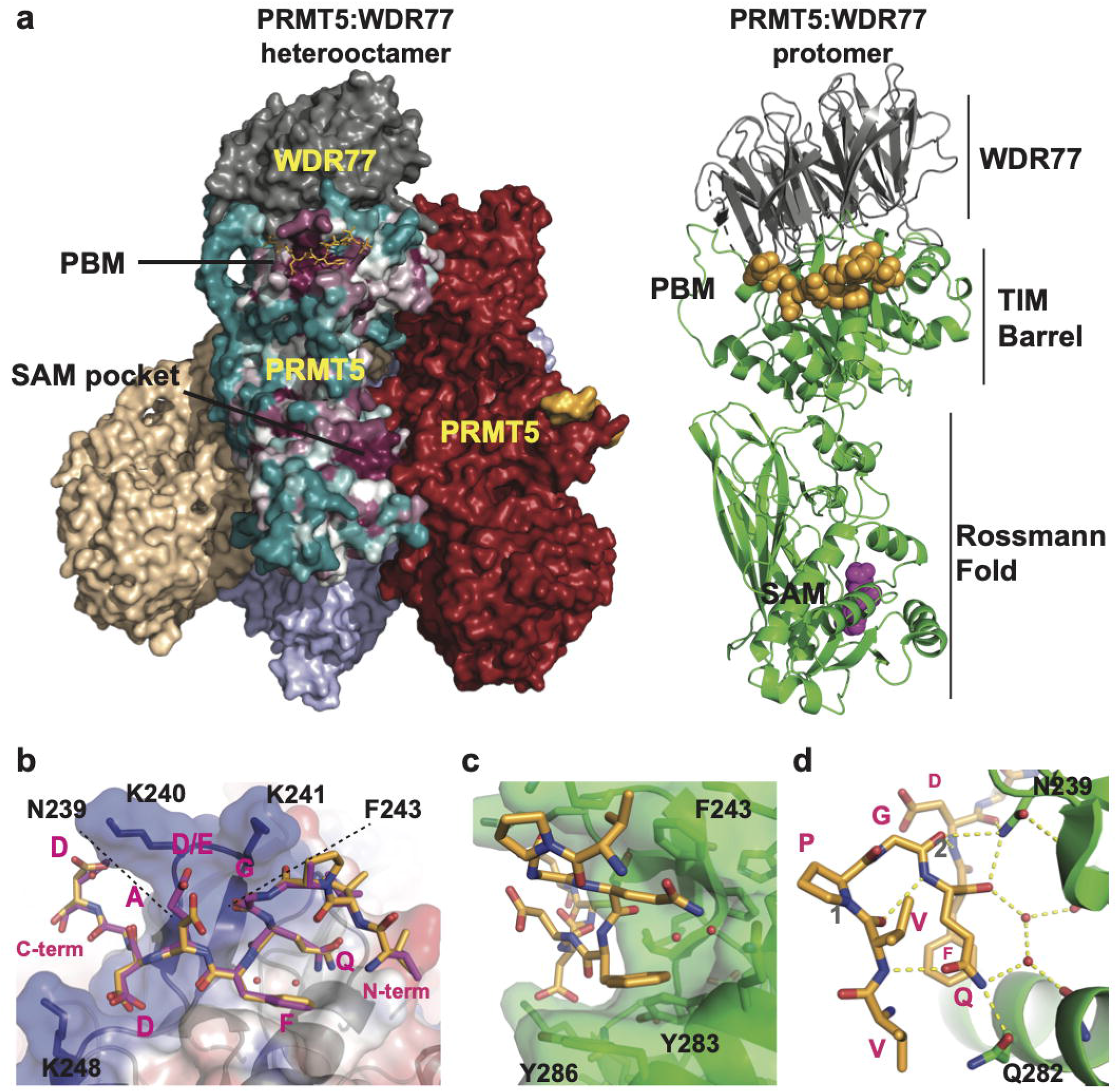
Structural basis for recognition of the PRMT5-binding motif (PBM) a. The PRMT5:WDR77 complex bound to the RIOK1 PBM peptide. Left, the hetero-octamer complex is shown as a surface with a different color for each protomer unit. The PRMT5 surface for the “front” protomer is colored by amino acid conservation on a sliding scale from cyan (least conserved) to maroon (most conserved). The RIOK1 PBM peptide is colored orange and shown as a stick representation (front protomer) or surface (all other protomers). Right, a single protomer unit of the complex is represented in cartoon form with PRMT5 in green and WDR77 in grey. The RIOK1 peptide and SAM are shown in orange and purple, respectively. b. PRMT5 (from the RIOK1 bound structure) is represented in cartoon form with a transparent surface colored by electrostatic potential from negative (red) to positive (blue). PBM peptides from RIOK1 (orange) and pICln (purple) are represented as sticks. Structures were aligned by minimization of global RMSD using Pymol. c. The PBM binding groove of PRMT5 (green) is shown as a cartoon with transparent surface. Two ordered water molecules are represented as red spheres. The RIOK1 peptide is shown in orange. d. Key hydrogen bonds of the RIOK1:PRMT5 interaction are shown as yellow dotted lines. Hydrogen bonds representing i+3 bonding pattern for β-turns labeled as “1” or “2” and described in the main text.

The interaction between the PBM peptide and PRMT5 occurs within a shallow groove of the TIM barrel, that is created by a protruding “finger” of two short antiparallel β-strands (Figure 2B). At the tip of the finger are two projecting lysine residues, K240 and K241. At the base of the finger are F243, which forms the back wall of the groove and N239, which serves several intermolecular hydrogen bonds (Figure 2C, 2D). A detailed view of this interaction is shown for the higher resolution RIOK1 structure (Fig. 2B-2D and S1).

The three main elements of PBM interaction are electrostatic attraction (Figure 2B), hydrophobic interaction of the peptide phenylalanine (Figure 2C), and a complex hydrogen-bonding network (Figure 2D). Both RIOK1 and pICln peptides have a strong overall negative charge with 3 ordered acidic side-chains in proximity to and complemented by three conserved lysine residues in the PRMT5 binding site (Figure 2B). In addition, the PRMT5 binding site has a shallow, hydrophobic “pocket”, the floor of which is composed of Y283 and Y286 while the back wall is formed by F243 of the β-strand finger (Figure 2C). Within this pocket are the phenyl ring and the β/γ carbons of the peptide Phe and Gln side-chains, respectively (Figure 2C). The positioning of these side-chains is enforced by an unusual peptide conformation discussed below. Also contained within the pocket are two ordered waters.

The PBM-PRMT5 interaction is promoted by several intermolecular and intrapeptide hydrogen bonds (Figure 2D). PRMT5 N239 has a prominent role in the interaction network and bonds with the main-chain carbonyl groups of 3 peptide residues. Two water molecules also bridge the main-chain of PRMT5 via hydrogen bonds to the peptide Gln. Notably, the positioning of the peptide Gln and Phe side-chains is enabled by an unusual peptide conformation requiring two consecutive β-turn motifs. The first turn is facilitated by the RIOK1 Pro-Gly sequence (Ala-Gly in pICln or Thr-Gly in COPR5) with typical i+3 bonding pattern (“1” in Figure 2D), but with the addition of a second stabilizing main-chain (valine amide) to side-chain (glutamine ε-carbonyl) bond. The second β-turn (“2” in Figure 2D) forces the Gln and Phe residues into the “stacked” conformation required for interaction with the shallow pocket described above.

### The PBM is necessary and sufficient for substrate adaptor binding to the methylosome

To determine whether the PBM is necessary for full-length adaptor protein binding to PRMT5, SPR was performed using the PRMT5:WDR77 complex and pICln or pICln lacking the PBM (pICln^ΔPBM^: deletion of residues 228-237). Concentration-dependent binding was observed for full length pICln, but not pICln^ΔPBM^ (Fig. 3A).

**Figure 3.**
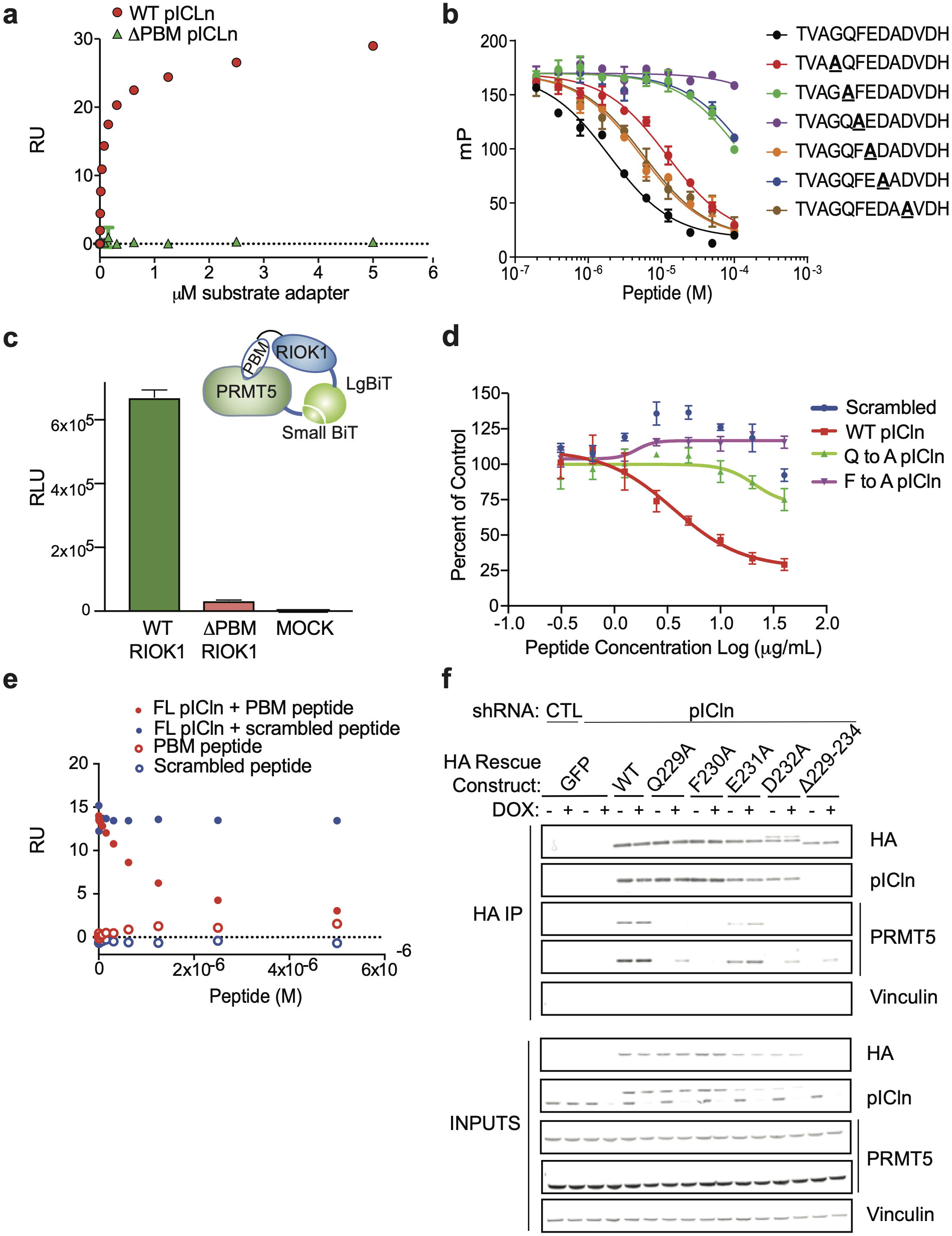
The PBM is required for PRMT5 binding to substrate adaptor proteins. a. Interactions between the PRMT5 complex and pICln protein were measured by SPR. The PRMT5 complex was immobilized by biotin-neutravidin and then full-length or mutant pICln protein was titrated as analyte. The RU response at equilibrium is shown. b. Competitive fluorescence polarization (FP) assay using a fluorescent RIOK1 peptide as probe. The indicated peptides were titrated as competitor molecules. c. The relative interaction between PRMT5 and RIOK1 or RIOK1^ΔPBM^ was measured in cells via NanoBiT assay in HEK293T cells following transient transfection. The N-termini of RIOK1 and PRMT5 were tagged with complementary “LargeBiT” and “SmallBiT” sequences, respectively. Relative luciferase signal was assayed as a measure of functional complementation. d. The NanoBiT PRMT5:RIOK1 system was used to test the ability of PBM peptides to compete for PRMT5 binding. Prior to addition of peptides, cells were permeabilized with 0.05% NP-40. Peptides of WT or point mutant PBM sequence were titrated into the permeabilized cells as indicated. Relative luciferase signal was assayed as a measure of functional complementation. e. Competition between full-length pICln and PBM peptide as measured by SPR. The PRMT5 complex was immobilized by biotin-neutravidin interaction. 125 nM pICln was co-injected with increasing concentrations of the indicated peptides. For comparison, the RU response with peptide alone is also shown. The RU response at equilibrium is shown. f. Co-immunoprecipitation of HA-tagged WT or PBM mutant pICln in the presence or absence of endogenous pICln depletion (dox inducible shRNA). Immunoblot detects the amounts of immunoprecipitated PRMT5 and pICln. Two exposures of PRMT5 are shown to visualize the dynamics of the binding between pICln^WT^ and pICln mutants. Vinculin serves as a loading control.

To ascertain which residues of the PBM are critical for the *in vitro* PRMT5:substrate adaptor interaction, fluorescence polarization (FP) was used to assess binding between the PRMT5:WDR77 complex and a fluorescently labeled PBM peptide from RIOK1. Unlabeled wild-type or single alanine mutant versions of the pICln peptide were then used as competitors. Mutation of any residue to alanine (except the existing Ala, not tested) within the consensus PBM had a deleterious effect on its ability to compete with the fluorescent peptide. The largest effects were observed for mutations corresponding to pICln^F230A^ followed by pICln^Q229A^ and pICln^D232A^ (numbering based on pICln; Figure 3B).

To examine whether the PBM is required for cellular interaction, HEK293T cells were stably infected with a PRMT5^WT^ Small-BiT fusion protein and a Large-BiT fusion to either RIOK1^WT^ or a ΔPBM mutant lacking residues 2-23 (RIOK1^ΔPBM^). Productive interaction between PRMT5-SmBiT and RIOK1-LgBiT is expected to reconstitute luciferase activity (Fig. 3C)^32^. Deletion of the PBM sequence led to a ~98% decrease in luciferase activity relative to WT RIOK1, suggesting that the PBM sequence is required for PRMT5:RIOK1 interactions in intact cells (Fig 3C).

To test whether the PBM peptide was *sufficient* to disrupt this NanoBiT interaction, cells were permeabilized and incubated with competitor peptide. pICln^WT^ PBM peptide efficiently displaced RIOK1-LgBiT from PRMT5 (IC_50_= 1.7 μM; Figure 3D). In contrast, the pICln^Q229A^ mutant peptide partially disrupted binding at high concentrations while the pICln^F230A^ mutant peptide failed to compete with RIOK1: PRMT5 binding (Figure 3D). Next, using SPR we tested the ability of this PBM peptide to compete with full-length pICln for PRMT5 binding (Figure 3E). A dose-dependent reduction in pICln binding was observed for PBM peptide, but not for a scrambled control peptide, consistent with the notion that disruption of the PBM interaction is sufficient for disruption of the full-length protein:protein interaction (Figure 3E).

To assess which PBM residues are critical for PRMT5:pICln complex formation in cells, co-immunoprecipitations (co-IP) of HA-tagged pICln or the indicated mutants were performed with and without shRNA knockdown of endogenous pICln (Fig. 3F). Relative to pICln^WT^, each pICln PBM mutant co-precipitated substantially less PRMT5 (Fig. 3F). Where endogenous pICln is also present (-dox), mutations of the PBM (except E231A) completely abrogates PRMT5 binding; where endogenous pICln is depleted (+dox), weak binding is still detected between the pICln PBM mutants and PRMT5 (Fig. 3F). Consistent with *in vitro* competition experiments (Fig. 3B,D) among the alanine mutants, pICln^F230A^ was the most impaired (Fig. 3F). Together, these data demonstrate the PBM peptide is necessary for full engagement of the pICln and RIOK1 substrate adaptors to PRMT5.

### The PBM groove is necessary for substrate adaptor binding to the PRMT5 methylosome

Next, to assess the importance of the complementary PRMT5 surface groove for substrate adaptor binding, structure-guided single point mutants and a composite, triple point mutant of HA-tagged PRMT5 (N239A, K240D, F243A), termed PRMT5^ADA^, were produced. The HA-tagged constructs were expressed with and without shRNA knockdown of endogenous PRMT5 as endogenous and exogenous molecules might co-assemble in the same hetero-octamer. Immunoprecipitates of HA-tagged PRMT5 proteins were assessed for co-purification of pICln by immunoblot (Figure 4A). In the absence of endogenous PRMT5 knockdown, the PRMT5^K240D^ and PRMT5^F243A^ single mutants showed partial loss of binding compared to wild-type PRMT5 (PRMT5^WT^), while the PRMT5^ADA^ triple mutant and PRMT5^N239A^ mutant showed nearly complete loss of pICln binding (Figure 4A). With knockdown of endogenous PRMT5, we did not detect co-precipitated pICln with any of the mutants (Figure 4A).

**Figure 4.**
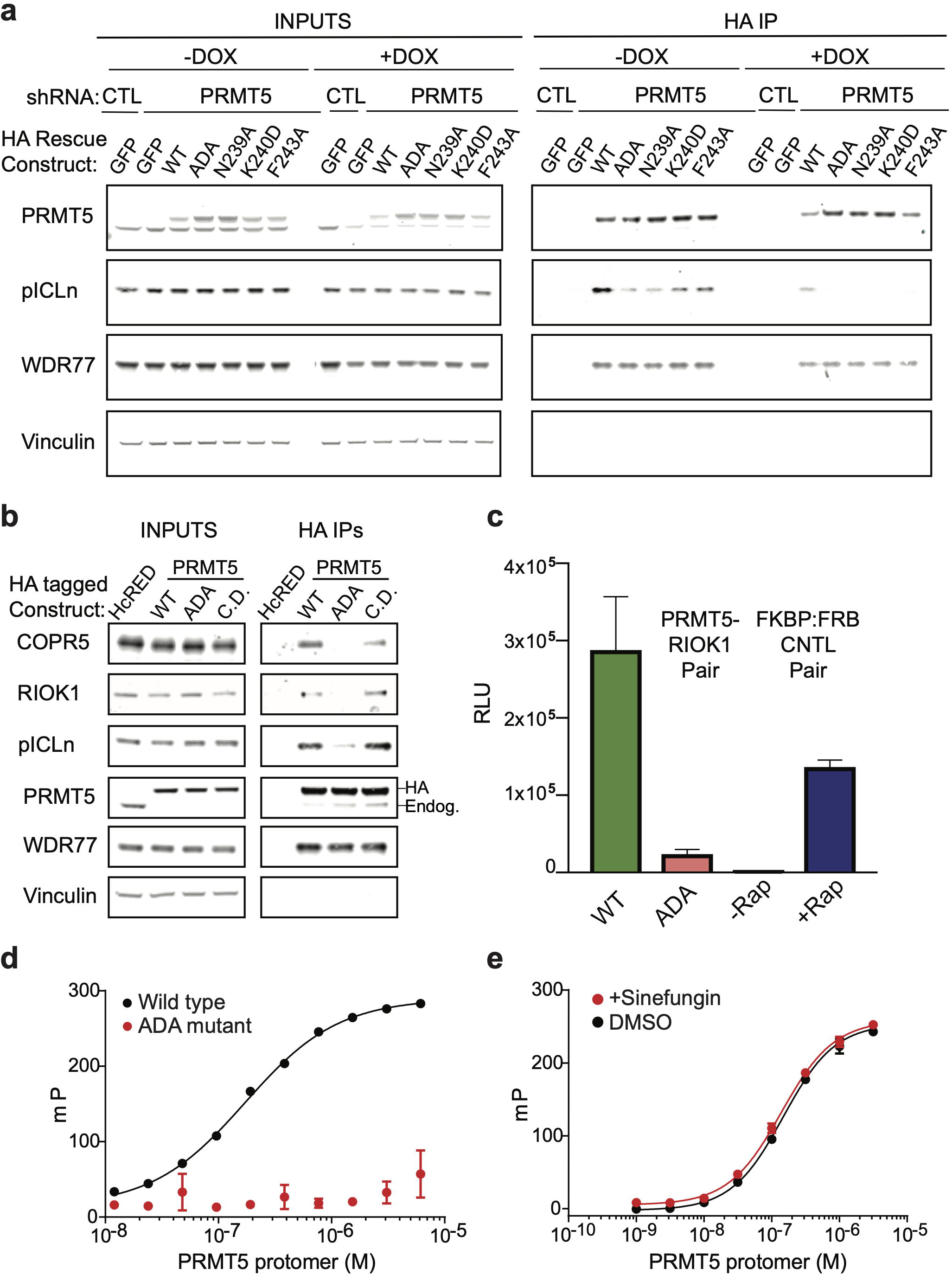
The PBM groove is required for PRMT5 binding to substrate adaptor proteins. a. Co-immunoprecipitation of HA tagged WT or PBM groove mutant PRMT5 in the presence or absence of endogenous PRMT5 depletion (dox. inducible shRNA) was performed. Immunoblot detects amounts of immunoprecipitated PRMT5, WDR77 and pICln. Vinculin serves as a loading control. b. Co-immunoprecipitation of HA tagged WT, PBM groove mutant (ADA), or catalytically dead (CD) PRMT5 in the presence or absence of endogenous PRMT5 depletion (CRISPr Cas9 KO). Immunoblot detects the amount of immunoprecipitated PRMT5, WDR77, pICln, RIOK1, and COPR5. Vinculin serves as a loading control. c. Mutations of the PBM interface on PRMT5 were tested via cell-based NanoBiT as previously described. PRMT5^WT^ or PRMT5^ADA^ mutant (N239A/K240D/F243A) and RIOK1 were tagged with complementary NanoBiT sequences and co-expressed. For comparison, NanoBiT complementation between the manufacturers’ rapamycin inducible control pair (FKBP-FRB; +/- Rap.) is also shown. Relative luciferase signal was assayed as a measure of functional complementation. d. Polarization of a fluorescent PBM peptide derived from RIOK1 was measured across the indicated concentrations of either WT or ADA PRMT5. e. Polarization of a fluorescent PBM peptide derived from RIOK1 was measured across the indicated concentrations of PRMT5 co-incubated with or without SAM analog sinefungin.

Next, binding of RIOK1 and COPR5 substrate adaptors was examined by expressing HA-tagged PRMT5 as either WT, ADA, or catalytically dead (“PRMT5^CD^”, R368A^24^) forms and using CRISPR-Cas9 to deplete endogenous PRMT5. HA-PRMT5 proteins were then immunoprecipitated and probed for copurification of pICln, RIOK1 or COPR5 by immunoblot. All three substrate adaptors were detected in immunoprecipitates of the WT or CD forms of PRMT5 while PRMT5^ADA^ exhibited 85-95% loss of substrate adaptor binding compared to control (Figure 4B).

To evaluate complex formation in live cells, the PRMT5:RIOK1 NanoBiT complementation assay was utilized where RIOK1-LgBiT was co-expressed with either WT or ADA mutant forms of PRMT5-SmBiT. As a control, a rapamycin-inducible complex between FKBP-SmBiT and FRB-LgBiT was included. Here, PRMT5^ADA^ showed significantly reduced binding compared to WT, suggesting that the PRMT5 groove is required for full RIOK1 adaptor binding (Figure 4C: 92% reduction with PRMT^ADA^ versus PRMT5^WT^).

We next assessed the effect of the ADA groove mutation using purified components *in vitro.* PRMT5:WDR77 and PRMT5^ADA^:WDR77 complexes were purified and evaluated for interaction with a fluorescently-tagged PBM peptide derived from RIOK1, as previously described. As determined by FP, PRMT5^WT^ interacted with the PBM peptide with an apparent K_D_=172 nM (Figure 4D). In contrast, a dissociation constant between PRMT5^ADA^ and the PBM peptide could not be determined (highest concentration tested = 50 μM PRMT5). Taken together these data suggest that the PBM groove of PRMT5 is necessary for the binding of pICln, RIOK1 and COPR5 substrate adaptors to the PRMT5:WDR77 methylosome.

### The PRMT5 Binding Motif binds independently of occupancy of the catalytic site

To determine whether the PBM interaction is independent of the catalytic site, PBM peptide binding to PRMT5:WDR77 complex was tested in the presence or absence of 30 μM sinefungin, a naturally occurring SAM analog and competitive inhibitor of PRMT5. As measured by FP, PBM peptide binding to PRMT5 was unaffected by sinefungin (Figure 4E). Similarly, no change in substrate adaptor binding was observed in co-IPs of HA-tagged PRMT5 after 24 hours treatment of cells with increasing concentrations of MTA, a metabolite inhibitor of PRMT5 (Figure S2). In agreement with biophysical and X-ray data, PBM binding to PRMT5 was unaffected by sinefungin or MTA, and hence is independent of catalytic site inhibition/occupation.

### The PRMT5 Binding Motif-PBM groove interaction is necessary for substrate methylation

To examine whether the PBM interaction is required for substrate methylation, PRMT5^WT^ and binding deficient mutants were expressed in the context of inactivation of endogenous PRMT5. Specifically, MiaPaca2 or HCT116 cells stably expressing Cas9 were infected with lentiviruses bearing sgRNA resistant cDNAs of HA-tagged PRMT5^WT^ or PRMT5 mutants and then infected with sgRNA targeting endogenous PRMT5. PRMT5 knockout efficiency was ~85% (Figure 5A). Levels of symmetric dimethylarginine (SDMA) in the presence of PRMT5^WT^, PRMT5^ADA^, and catalytically dead PRMT5 (PRMT5^CD^) mutants were assessed by immunoblot. Rescue expression of the control HcRed or of PRMT5^CD^ resulted in a substantial loss of SDMA, while expression of PRMT5^WT^ restored endogenous levels of SDMA. In contrast, rescue with the PBM-binding mutant (PRMT5^ADA^) affected a distinct subset of SDMA-modified proteins (Figure 5A; HCT116 cells were engineered to be MTAP-/-, described in Figure S4A). For the shared bands, the extent of methylation loss was similar between the PRMT5^CD^ and PRMT5^ADA^ mutant cells. These results suggested that there are PBM dependent and independent substrates of the PRMT5 methylosome (Figure 5A).

**Figure 5.**
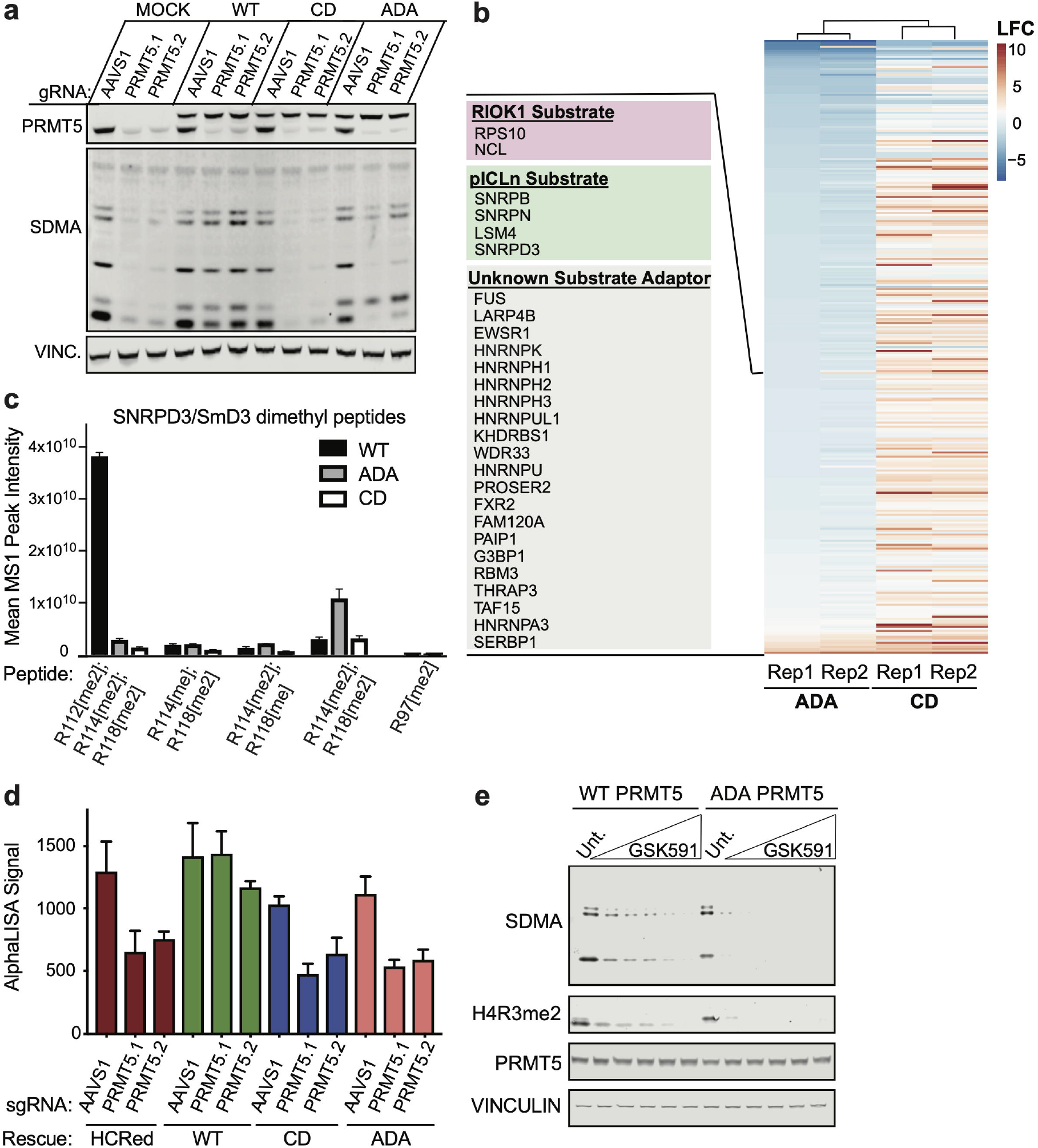
The PBM-PBM groove interaction is required for methylation of a subset of PRMT5 substrates. a. Symmetric dimethyl arginine (SDMA) detected by immunoblot in a PRMT5 KO-rescue system utilizing the PBM groove mutant (ADA), catalytically dead (CD) or wildtype (WT) PRMT5 in HCT116 MTAP KO cells. b. SDMA-modified peptides were immunoprecipitated from CD, ADA, and WT PRMT5 rescued MiaPaca2 cells following KO of endogenous PRMT5, and quantified by LC-MS/MS. Each row represents a distinct SDMA peptide detected. Peptides are clustered by ADA Replicate 1. Peptides with a 2-fold decrease in ADA versus WT are labeled; list is in order of the methyl peptide of that protein with the largest FC except RIOK1 and pICLn substrates which were isolated for visual clarity. Full data - Table S2. c. The mean MS1 peak intensity for all dimethylated peptides of SmD3 that were detected by LC-MS/MS in (b) are shown for each the WT, ADA and CD PRMT5 cell lines. Peptides are ordered 1-5 based on their log fold-change WT/CD ~strength as a PRMT5 substrate. d. The relative amount of SDMA-modified SmB protein was measured by AlphaLISA proximity assay, as described in methods, for each HCT116 MTAP-/-: HcRed, WT, ADA and CD-PRMT5 expressing stable cell lines in the presence or absence of 2 PRMT5 targeting sgRNAs. e. The amount of total SDMA and H4R3 SDMA was measured by immunoblotting in the HCT116 MTAP-/-; endogenous PRMT5-/-; HA-PRMT5^WT^ or HA-PRMT5ADA cell lines following a titration of GSK591 (PRMT5 inhibitor) treatment for 96 hours. Doses:0-5-10-20-50-100nM.

To identify PRMT5 substrates that require the PBM interaction, SDMA immunoprecipitation mass spectrometry was performed (Figure 5B-C; Table S2). Protein lysates were prepared from the aforementioned PRMT5 KO-rescue MiaPaca2 cells expressing either PRMT5^WT^, PRMT5^ADA^, or PRMT5^CD^. Symmetrically dimethylated tryptic peptides were enriched by SDMA-IP and analyzed by LC-MS/MS. In total, we identified 408 unique dimethyl peptides from 107 proteins. Sixty-five of these SDMA-modified proteins were determined to be PRMT5 substrates (SDMA Peptides FC>2 CD/WT and SDMA event within “GRG” motif). Twenty-eight out of 65 total SDMA proteins from our analysis were also identified in a similar SDMA substrate LC-MS/MS study using HeLa cells (Table S2)^33^. Furthermore, 307 of the 408 SDMA peptides identified in our study share a common “GRG” sequence consistent with a structural requirement for this motif to interact with the PRMT5 catalytic site and has been reported to be the primary PRMT5 substrate motif^34,35^. Thus, we are able to robustly detect SDMA events and PRMT5 methylation sites.

Twenty-five of the 65 PRMT5 substrates detected herein appeared to be PBM-dependent (Figure 5B; SDMA Peptides FC>2 compared to WT). Consistent with the immunoblot results, a subset of PRMT5 catalytic substrates were PBM independent (Figure 5A-B; immunoblots of PRMT5 and SDMA levels in MiaPaca2 cells shown in Figure S3A). The PBM-dependent substrates, including known RIOK1-dependent substrates (nucleolin/NCL and RPS10) and known pICln-dependent substrates (SmB, SmD3, SMN and LSM4), were all markedly decreased in cells expressing the PRMT5^ADA^ mutant. These results were consistent with a requirement of the PBM interaction for their methylation^12,14–16^(Figure 5B-C). Histones were largely not detected in this study likely due to their Arg and Lys rich nature not being compatible with tryptic digest. A number of substrates were also identified as PBM groove-dependent for which no substrate adaptor has been identified, such as SERBP, which had the strongest SDMA peptide fold change in PRMT5^ADA^ compared to PRMT5^WT^ (Figure 5B). Pathway enrichment analysis demonstrated that both PRMT5ADA and PRMT5^WT^ substrates are primarily from RNA binding and RNA processing pathways. While 38% of PRMT5 substrates identified here were found to be PBM-dependent, no differences were determined between pathways regulated by PRMT5ADA versus PRMT5^WT^ (Figure S3B). These data are in-agreement with reported roles of PRMT5 in the regulation of RNA biology^17,34^. Occasional enhancement of dimethylated peptides was observed in the PRMT5^ADA^ rescue cells; however, these appear to be partially methylated forms of substrate peptides that were otherwise decreased. For example, SmD3 dimethyl R114, R118 increases concomitant with a significant decrease in R112, R114, R118 tri-dimethylated peptide (Figure 5C), suggesting that the PBM binding site is required for SmD3 methylation overall and that the R112 may be the most PBM groove-dependent of these 3 sites.

To further test the requirement of the PRMT5 PBM groove for the known pICln-dependent methylation of Sm proteins, an AlphaLISA proximity assay was developed using antibodies against SmB (Gene name: SNRPB) and SDMA^32^. Using the PRMT5 KO-rescue system, the SDMA-SmB proximity signal decreased in both the PRMT5^ADA^ and PRMT5^CD^ mutants compared to PRMT5^WT^ (Figure 5D). To examine whether loss of PBM binding enhances the loss of methylation induced by PRMT5 catalytic inhibitors, the aforementioned PRMT5^WT^ and PRMT5^ADA^ cells were treated with 5-100nM of GSK591 and total SDMA and H4R3me2 (a known COPR5-dependent PRMT5 substrate)^20^ levels determined by immunoblot after 96 hours (Figure 5F; these cells were single cell clones of endogenous PRMT5 KO, shown in Figure S4B)^33^. Compared to PRMT5^WT^ controls, cells containing PRMT5^ADA^ were sensitized to GSK591 and exhibited a substantial shift in inhibitor IC_50_ for both total SDMA and H4R3me2 (Figure 5E). Together these data suggest that the PBM groove plays a crucial role in substrate adaptor mediated substrate methylation.

### Mutation of the PBM binding groove impairs viability of MTAP null cell lines

To assess the requirement of the PBM groove for growth of *MTAP* null cells, a PRMT5 KO-rescue system was engineered in a CDKN2A-/-, MTAP-/- cell line (MiaPaca2) as well as in two isogenic pairs where *MTAP* was deleted using CRISPR-Cas9 (HCT116 and PK-1). In addition to rescue with PRMT5^WT^ and PRMT5^ADA^, a PRMT5CD construct served as a loss of function control. In all three MTAP null cell lines, ectopic PRMT5^WT^ rescued growth following depletion of the endogenous PRMT5 (Figure 6A-C, AAVS1 Incucyte data not shown). The amount of overexpression and efficiency of knockout were measured by immunoblotting at day 5 after infection with sgRNAs (Figure 6D); PRMT5 protein levels were diminished >85% in all cell lines. Cell lines expressing PRMT5^ADA^ showed a statistically significant hypomorphic growth phenotype (MiaPaca2: n=5, p<0.05) relative to cells expressing PRMT5^WT^.

**Figure 6.**
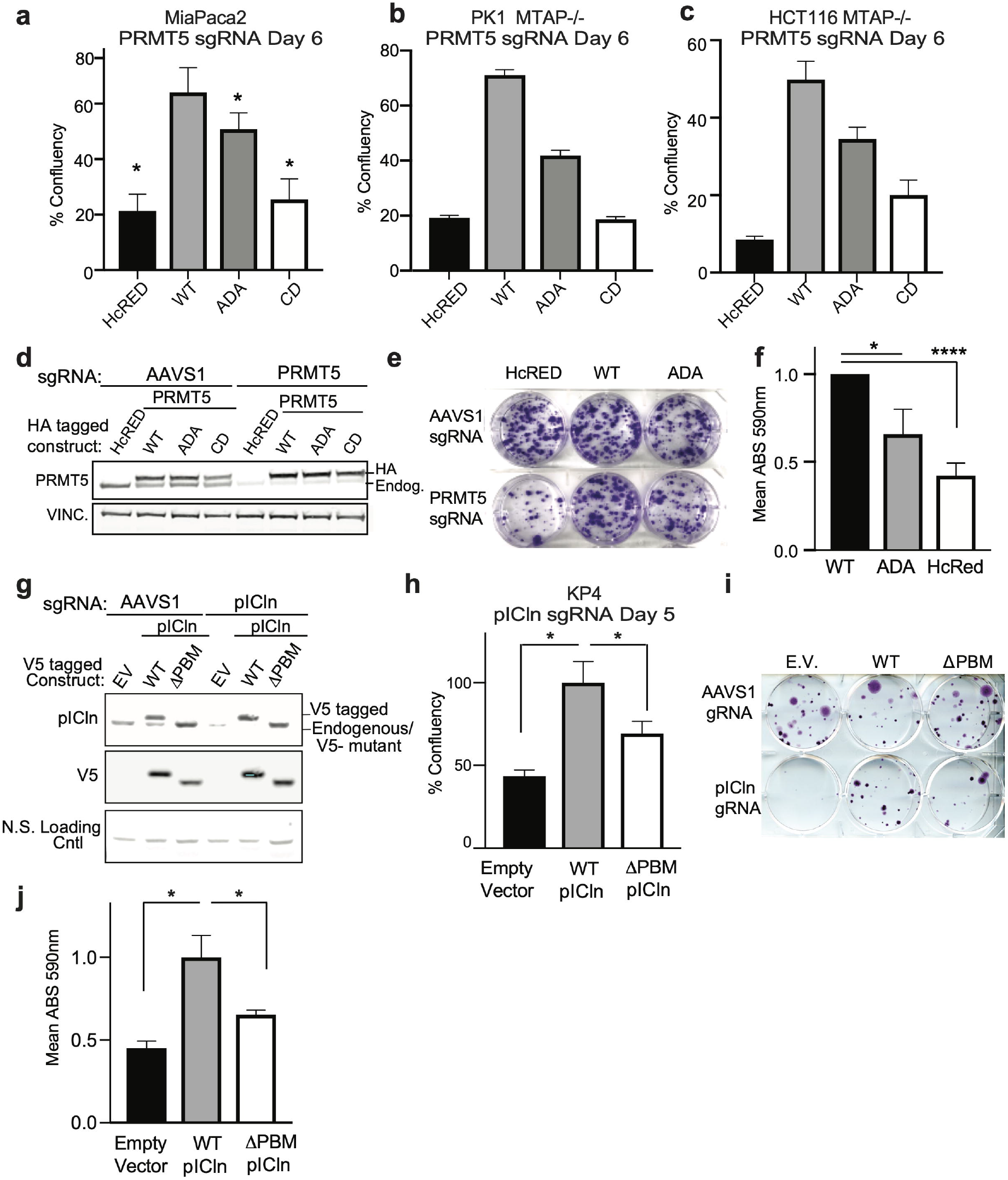
Mutation of the PBM interaction site impair growth in MTAP null cells. a. Incucyte cell growth plots of MiaPaca2 Cas9 expressing cells infected with PRMT5-targeting guide RNA to KO endogenous PRMT5 are shown. Cell lines are stably expressing HA-tagged HcRed, PRMT5^WT^, PRMT5ADA or PRMT5CD. 5 days after KO and 2 days after selection with puromycin, cells are seeded into growth assays. Growth was measured in the presence of 10μM MTA. Day 6 (T=156h) is plotted. Mean +/- SEM are shown for n=5 replicates; *=p<0.05 paired student’s t-test. b. Incucyte cell growth plots of PK1 MTAP-/-Cas9 expressing cells infected with PRMT5-targeting guide RNA to KO endogenous PRMT5. Cell lines are stably expressing HA-tagged HcRed, PRMT5^WT^, PRMT5ADA or PRMT5CD. Growth was measured in the presence of 10μM MTA. Day 6 (T=156h) is plotted; n=1. c. Incucyte cell growth plots of HCT116 MTAP-/- Cas9 expressing cells infected with PRMT5-targeting guide RNA to KO endogenous PRMT5. Cell lines are stably expressing HA-tagged HcRed, PRMT5^WT^, PRMT5ADA or PRMT5CD. Growth was measured in the presence of 10μM MTA. Day 6 (T=152h) is plotted; One representative experiment is shown, n=2 replicates in HCT116 cell line, n=8 total replicates across 3 cell lines in (A-C). d. Immunoblots of the genotypes from (A) MiaPaca2 cells. Immunoblotting was used to detect endogenous and HA-tagged PRMT5 in each cell line: HcRed, PRMT5^WT^, PRMT5ADA or PRMT5CD in the presence of AAVS1 control or PRMT5 targeting sgRNA. e. Crystal violet-stained colony formation assay of MiaPaca2 Cas9 expressing cells infected with AAVS1 control or PRMT5-targeting guide RNA to KO endogenous PRMT5 and rescue with either HcRed, PRMT5^WT^, or PRMT5ADA in the presence of 10μM MTA; 14 days. One representative experiment is shown, n=5. f. Quantification of colony formation assays in (E) using acid-solubilization and ABS reading. Normalized mean ABS is shown +/- SEM of n=5 experiments. *=p<0.05 student’s t-test; ****=p<0.0001 student’s t-test. g. Immunoblots of KP4 Cas9 expressing cells stably expressing V5-tagged pICln and EV constructs after infection with pICln targeting sgRNA or control (AAVS1) sgRNA. Anti-pICln and anti-V5 measure knockout and overexpression efficiency. h. Incucyte cell growth plots of KP4 Cas9 expressing cells infected with pICln-targeting guide RNA to KO endogenous pICln are shown. 5 days after KO and 2 days after selection with puromycin, cells are seeded into growth assays. Cell lines are stably expressing Empty Vector, pICln^WT^, or pICln^ΔPBM^ truncation mutant. EV vs WT p=0.02; pICln^WT^ vs pICln^ΔPBM^ p=0.04; n=4 replicates. i. Colony formation assay stained with crystal violet of KP4 Cas9 expressing cells infected with AAVS1 control or pICln-targeting guide RNA to KO endogenous pICln and rescue with either Empty Vector, pICln^WT^, or pICln^ΔPBM^ truncation mutant. 5 days after KO and 2 days after selection with puromycin, cells are seeded into growth assays. j. Quantification and statistical analysis of the biological replicates in (I) (EV vs pICln^WT^ p=0.01; pICln^WT^ vs pICln^ΔPBM^ p=0.04; n=3).

To test the requirement of the PBM groove in an orthogonal assay system and for longer term growth, colony formation assays were performed using MiaPaca2 cells. Knockout of endogenous PRMT5 showed a substantial reduction in colony number (HcRED mock rescue) that was rescued by re-introduction of PRMT5^WT^, consistent with an important role of PRMT5 function in colony growth formation. Rescue with PRMT5^ADA^ again showed a hypomorphic growth phenotype with a decrease in colony number relative to PRMT5^WT^ (n=5, p<0.05), (Figure 6E-F). Taken together, these data suggest that perturbation of the PBM groove on PRMT5 confers a hypomorphic loss of cell viability.

### Mutation of the pICln PBM impairs viability of MTAP null cells

In addition to PRMT5, both shRNA knockdown of pICln and RIOK1 are context specific dependencies in *MTAP-/-* lines and are common essential genes when targeted by sgRNAs^1,22, 23^. To determine whether the pICln PBM is necessary for growth in an *MTAP* null line, KP4 pancreatic cancer cells, harboring a *CDKN2A, MTAP* co-deletion, and stably expressing CAS9 were infected with lentiviruses encoding either sgRNA-resistant pICln^WT^ or pICln^ΔPBM^ or an empty vector. The three resulting stable cell lines were then stably infected with lentivirus directing the expression of sgRNAs targeting either the AAVS1 safe-harbor site or pICln.

Knockout of pICln led to significantly reduced, but incomplete, loss of protein in pooled cells (Fig. 6G) and diminished growth in proliferation and colony forming assays (Fig. 6H-J) (colony formation WT:ΔPBM p=0.043; incucyte at day 5 WT:ΔPBM p=0.04, n=4). Growth defects were rescued by expression of V5-pICln^WT^, but growth rescue effects were attenuated when expressing V5-pICln^ΔPBM^. Specifically, the cells expressing pICln^ΔPBM^ showed a hypomorphic growth rescue when compared to the empty vector (~65% versus 35% rescued growth; Fig. 6H-J). Overall, these results suggest a hypomorphic decrease in MTAP^-/-^ cell viability following genetic perturbation of the pICln PBM.

## Discussion

PRMT5 is a critical regulator of symmetric arginine dimethylation and regulates key cellular processes such as RNA splicing, ribosome biogenesis and chromatin dynamics. Herein, we report a novel mechanism of interaction between the PRMT5 methylosome and its adaptor proteins that mediates substrate methylation through a structurally resolved binding site on PRMT5 (PBM groove) and a newly defined substrate adaptor motif (the PBM; Fig. 7). Though structurally distinct, parallels can be drawn to the cullin RING E3 ligase family where substrate adaptors also act as exchangeable partner proteins to recruit specific sets of substrates to the same core enzyme complex^36^. The substrate sequence for PRMT5 methylation is fairly non-specific (RG or GRG at the site of methylation), and the use of substrate adaptors permits PRMT5 to act in a highly selective manner. Here, we identify the PBM site as a mechanism required for this broad regulatory function across otherwise divergent substrates.

**Figure 7.**
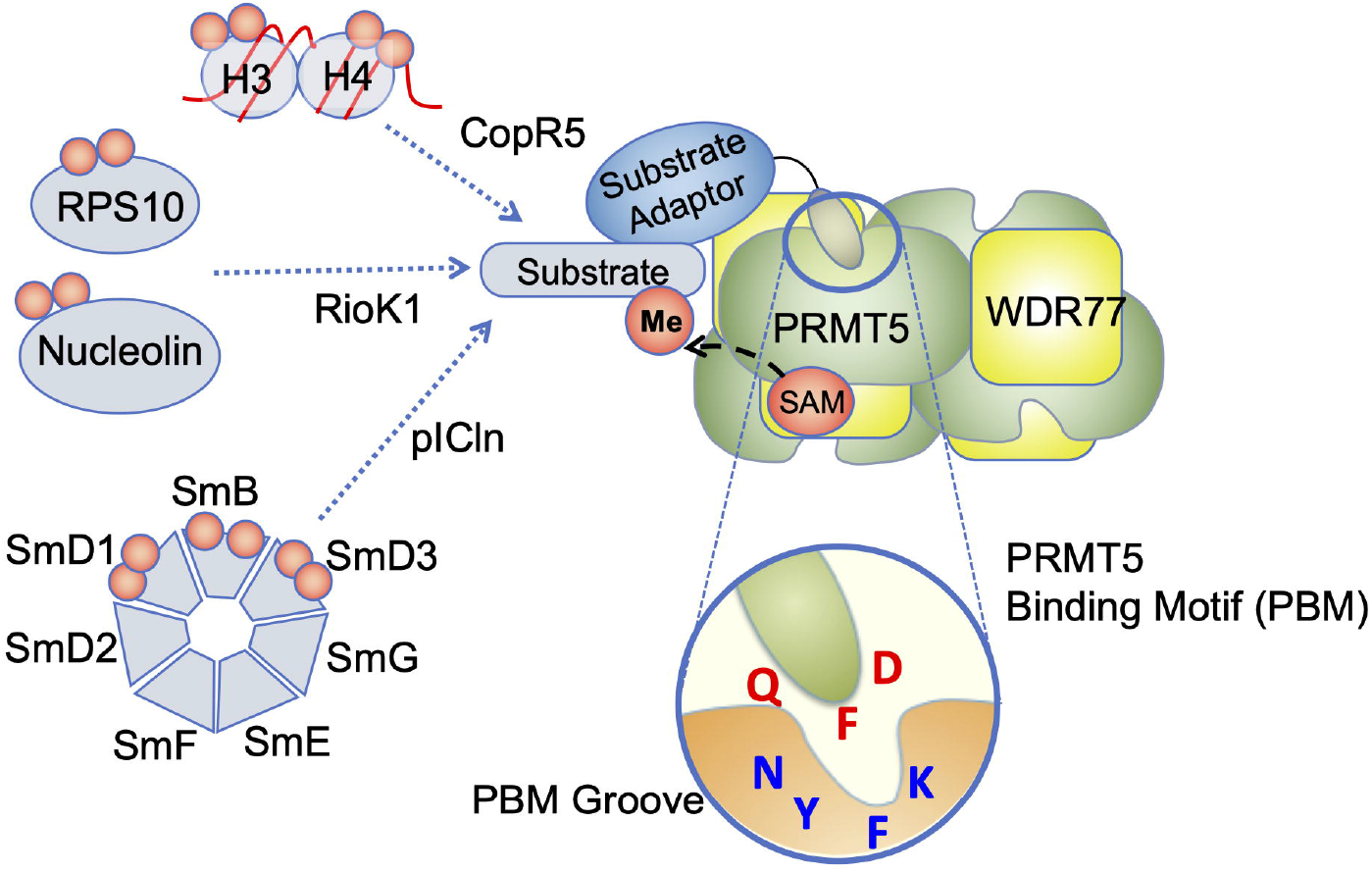
Graphical Model of PRMT5:substrate adaptor site and its cellular consequences. In Figure 7, we present a unified model for substrate recruitment to the PRMT5 enzyme. Substrate adaptor proteins function in an exchangeable, and mutually exclusive, manner via the PBM binding site. This allows for specific methylation and regulation of distinct substrate classes using the same core enzyme.

While the PRMT5ADA PBM groove mutant showed decreased methylation of many well-described PRMT5 substrates (Sm proteins, ribosomal proteins, and others), it did not lead to as strong of a viability phenotype compared to the catalytically dead “CD” mutant. There are multiple possibilities to explain this observed difference in cell viability. First, the greater effect of the PRMT5CD mutant could be due to loss of activity on one or more essential *PBM-independent* substrates. Second, it is possible there is a cumulative viability effect from loss of both PBM-dependent and independent substrates. A third possibility is that the PRMT5ADA and PRMT5CD mutations are same in kind but differ in degree of loss for some cell essential modification(s). The critical viability threshold for any specific methylation event is unknown. Lastly, it is possible that although the PRMT5ADA mutation significantly disrupts adaptor binding, as measured in multiple orthogonal assays, enough residual complex formation (as in Figure 4C) occurs to sustain a viability threshold. For example, if the substrate molecules have some level of adaptor-independent affinity for the PRMT5 complex at and proximal to their RG/GRG substrate motifs^9,31^, then this direct substrate contact could result in a level of independence from the adaptor:PRMT5 complex and subsequent methylation despite loss of the PBM component.

As PRMT5 is the target of numerous therapeutic efforts, it is important to identify substrates and cellular functions of PRMT5 and to differentiate between any substrates or functions unique to *MTAP/CDKN2A*-deleted cells as these may provide therapeutic vulnerabilities. The detection of PBM independent methylation events raises the possibility of substrate recruitment through other components of the methylosome. For example, 2 PRMT5 substrates for which no substrate adaptor has been identified, PRDM1 and ZNF326, share another unique 7-mer peptide sequence G[K/R]DLYRS (Figure 1B). Such a sequence would be highly unlikely to bind to the same PBM site based on the X-ray crystal structure, but could bind to a presently unidentified surface of the methylosome, such as the propeller domain of WDR77. Our current results do not identify the PBM as the exclusive mechanism for substrate recruitment, but indeed suggest there could be multiple paths to substrate specificity.

In conclusion, we have identified a novel means by which PRMT5 recruits substrates for methylation. The newly identified pocket of the methylosome is required for the binding of substrate adaptors pICln, RIOK1, and COPR5 and for the methylation of select PRMT5 substrates. Modulation of the PRMT5-substrate adaptor binding site leads to hypomorphic effects on cell growth in *MTAP* null cells. Thus, identification of the PBM site reveals a novel allosteric pocket for potential PRMT5 modulation. These results indicate that many PRMT5 substrates are modified by a two-step process: 1) recruitment via a PBM-adaptor and 2) catalysis of methyl transfer. To inhibit PRMT5 signaling in human diseases, one could envision blocking one or both of these steps. Given the effects of perturbing this site on cell growth coupled with previous findings that existing PRMT5 catalytic inhibitors are not selective for *MTAP* null cells^22^, this newly defined site on the PRMT5 methylosome may provide a novel therapeutic target in *CDKN2A/MTAP* deleted tumors.

## Methods

### Tissue Culture and cell lines

KP4, MiaPaca2, HCT116, PK1, and HEK293T cell lines were acquired from the Cancer Cell Line Encyclopedia (CCLE) at the Broad Institute^37,38^. These cell lines from the CCLE were acquired from ATCC and validated by short tandem repeat (STR) profiling analysis. All subsequent cell line derivatives underwent repeat STR analysis following lentivirus infection and selection. Cell lines were maintained in media (F12-DMEM for KP4 or DMEM for all other lines, Corning) supplemented with 10% fetal bovine serum (Sigma) and 1% penicillin/streptomycin (Gibco) in a humidified 37 C incubator at 5% CO_2_. Cells were not kept in culture beyond 15 passages.

### CRISPR-Cas9

Cas9 expressing cells were generated and characterized for Cas9 cutting efficiency as previously described^39^ (Genetic Perturbation Platform, Broad Institute). All cell lines utilized herein showed greater than 90% Cas9 efficiency in this assay. Guide RNAs (sgRNAs) were designed using the Genetics Perturbation Platform tool and either guides targeting intron-exon junctions were chosen or codon optimization was performed to allow selective knockout of endogenous genes while allowing expression of ectopic cDNAs. Guide RNAs were cloned into pXPR_003 or pXPR_043 by BsmBI digest. For MTAP and NTC, guide RNAs were cloned into pXPR_023 using BsmBI digest. pICln guide 2 was acquired from Cellecta in vector pRSG17. The following guide RNA sequences were used: PRMT5 gRNA 1: GGTTGCTACTCACGTCACCA; PRMT5 gRNA 2: ATACAGCTTTATCCGCCGGT; pICln gRNA 1: GTCACACGCTGATTTATCACT; pICln gRNA 2: GCCACACTGGAGAGATTAGA; AAVS1 gRNA: GGGGCCACTAGGGACAGGAT; NTC sgRNA: GTATCCTGACCTACGCGCTG; MTAP sgRNA: GGCTCATCTCACCTTCACGG.

### Generation of KO Clonal Pooled Cell Lines

HCT116 and PK1 were infected with MTAP sgRNA or NTC sgRNA control in pXPR023 Cas9 all in one vector, selected with blasticidin (2μg/mL) and seeded into 96 well plates at a limiting dilution, allowing less than 1 cell per well. After colonies were formed and allowed to grow for ~4 weeks, cell lysates were generated from each and assessed for MTAP protein expression. Three HCT116 clones and 3 PK1 clones were determined to have MTAP loss and were pooled for subsequent assays (HCT116 clones # 3, 8, 10; Figure S4B). HCT116 MTAP-/- and PK1 MTAP-/- cells above were then infected with HcRED, PRMT5^WT^, PRMT5ADA or PRMT5CD in pLX305 and selected with hygromycin. Finally, HCT116 MTAP-/- PRMT5^WT^ and PRMT5^ADA^ cell lines were infected with PRMT5 sgRNA 1 in BRD003 vector, selected with puromycin (1μg/mL) and seeded into 96 well plates at a limiting dilution, allowing less than 1 cell per well. Four PRMT5^WT^ and five PRMT5^ADA^ clones were determined to be PRMT5-/- and were subsequently pooled (PRMT5^WT^ clones #1, 5, 7, and 15 and PRMT5ADA clones #5, 6, 13, 14 and 17; Figure S4B).

### Inducible shRNA

For PRMT5 cellular binding assays, MiaPaca2 cells were infected with pLKO shRNA lentiviruses containing either a non-targeting control shRNA or PRMT5 targeting shRNAs. The sequences of the PRMT5 shRNAs were previously reported^22^. To perform pICln cellular binding assays, MiaPaca2 cells were infected with pLKO shRNA lentiviruses containing either a non-targeting control or 1 of 2 pICln targeting shRNAs. These had the following sequences: pICln shRNA 1: CCCTCTGAGTAGGCCTATAAT; pICln shRNA 2: GCCTAGTGATAAATCAGCGTT. These and all new constructs in the manuscript have been deposited to AddGene.

### Mammalian Expression Constructs

PRMT5 was cloned into pLX305 PGK 3X HA Hygro-2A-GFP vector using Gibson Assembly. Point mutations in PRMT5 were generated using the Q5 site directed mutagenesis kit (New England Biolabs). HcRed was subcloned from pDONR223 by Gateway cloning into pLX305 PGK 3X HA Hygro-2A-GFP. pICln was cloned into an N-terminally V5-tagged vector derived from the pLX307 backbone (pSL307). Deletions and point mutations were generated as above for PRMT5.

### Cell Lysis, immunoblots and immunoprecipitations

For HA or endogenous target immunoprecipitation (IP), cells were lysed in 0.1% NP-40 co-IP lysis buffer (10% glycerol, 50 mM Tris-HCl, 150 mM NaCl, 2 mM EDTA, 0.1% NP-40, supplemented with protease inhibitors (Thermo Fisher) and phosphatase inhibitor (Thermo Fisher) and passed through a 26 ½ gauge needle 3 times. Lysates were cleared by centrifugation, quantified by bicinchoninic assay (BCA; Thermo Fisher) and incubated with anti-HA resin (Sigma) rotating at 4C for 1h, washed with lysis buffer and eluted from the beads with a 1:1:1 mixture of 0.5M DTT; NuPAGE 4X SDS loading buffer (Life Technologies); co-IP lysis buffer and boiled for 10 minutes.

For total protein detection cells were lysed in RIPA buffer (1% NP-40, 0.1% SDS, 0.25% sodium deoxycholate, 150 mM NaCl, 10% glycerol, 25 mM Tris, 2mM EDTA) supplemented with protease inhibitors and phosphatase inhibitor. Protein lysates were quantified by BCA kit (Thermo Fisher) and 20-30μg of protein were loaded per lane. Lysates were resolved on 4-12% SDS-PAGE NuPage gradient gels (Invitrogen), transferred to nitrocellulose membranes, blocked using LiCor PBS blocking buffer and probed using the following antibodies: anti-PRMT5 (Abcam ab31751); anti-SDMA (Cell Signaling 13222); anti-HA (Abcam ab18181); anti-pICln/CLNS1A (Novus NBP2-33958 or Santa Cruz sc-271454); anti-SmB/B’/N (Santa Cruz sc-130670); anti-SmB/ SNRPB (Thermo Fisher MA513449); anti-SmD1 (Novus NBP2-36427); anti-Vinculin (Sigma V9131); anti-nucleolin (Abcam ab22758); anti-WDR77/MEP50 (Cell Signaling 2828) anti-RIOK1 (Bethyl A302-457A); anti-COPR5 (Novus NBP2-30884); anti-V5 (Cell Signaling 13202S); or anti-MTAP (Santa Cruz sc-100782) as indicated. Image quantification was performed using the LI-COR imaging suite (Image Studio Lite) where all blots were determined to be in the linear range by the software.

### NanoBiT Assays

Detection of protein interactions using split luciferase complementation assays were performed as follows. pFC34K LgBiT TK-neo, pFC36K SmBiT TK-neo, pFN33K LgBiT TK-neo and pFN35K SmBiT TK-neo Flexi Vectors were purchased (Promega). Gibson Assembly was performed to clone the cDNA sequence encoding the SmBit peptide sequence to the N-terminus of PRMT5 in pLX304-Blast and the cDNA sequence encoding the LgBit protein to the N-terminus of RIOK1 in pLentiCMV-Puro. HEK293T cells were co-infected with lentiviral pLX304 SmBit-PRMT5 and pLentiCMV LgBit-RIOK1 and selected with puromycin (1ug/mL) and blasticidin (3μg/ml). Mutations and truncations of RIOK1 and PRMT5 were introduced using Gibson Assembly. For transient assays, plasmids (30 ng each) were introduced by transient transfection using TransIT (Mirus Bio.). Specifically, HEK293T cells were plated in 96 well plates at 10,000 cells/well, and the following day were transfected with either SmBit-PRMT5^WT^ or PRMT5ADA mutant together with either LgBit-RIOK1^WT^ or LgBit-RIOK1^ΔPBM^ mutant. Luciferase assays were performed 24h after transient transfection. For peptide displacement assays, HEK293T cells stably expressing SmBit-PRMT5 and LgBit-RIOK1 were trypsinized, resuspended in serum-free media and diluted to 100,000-300,000 cells/well. Cells were permeabilized using 0.05% NP40 for 5 minutes at room temperature (RT). Peptides were added in increasing doses as indicated simultaneously with the addition of NP40. After 5 minutes, NanoLuc reagent was added and read immediately. In both transient and stable NanoLuc experiments, data were collected in black, 96 well tissue culture coated plates (Corning) using a Perkin-Elmer Envision plate reader.

### IncuCyte assays

Cell lines were seeded in white 96 well plates (Corning) and monitored for growth for 7-10 days on an IncuCyte (Essen BioSciences). KP4 cells-500 cells/ well; MiaPacs2 cells: 1,000 cells/well. All conditions were plated in triplicate and 4 images per well were collected every 4 hours. Growth curves were generated in the Incucyte software. Statistical analysis was performed in GraphPad Prism. For competition assays, cells were monitored in both fluorescence and phase fields of the IncuCyte.

### Colony Formation Assays

After determining cell counts using a ViaCell counter, cells were seeded in technical triplicates at a limiting density of 500-1,000 cells per well in a 6 well tissue culture coated dish (Corning) to allow single cell colony formation. Cells were allowed to grow for 10-21 days (until colonies were visible in the control wells). To determine colony numbers, the media was removed and cells were washed twice in DPBS, fixed in 10% formalin for 30 minutes, and stained with 2 mL crystal violet (0.05% crystal violet v/v in 10% methanol and water) for 1h at RT. Stain was removed, wells were washed 3 times in diH_2_O and dried overnight. Colonies were imaged in Photoshop and quantified by glacial acetic acid solubilization and dye intensity was measured by absorbance reading at 570 nm.

### AlphaLISA Assays

AlphaLISA assays were performed according to the manufacturer’s instructions (Perkin Elmer). Briefly, cells were lysed in AlphaLISA Immunoassay Buffer by rocking at 4°C for 1 hour. Lysates were cleared by centrifugation and proteins were quantified by BCA assay as above. 5 μg of protein was added to each well of a white 96 well plate (Corning). 3 nM anti-SmB/B’/N (clone 12F5, #130670, Santa Cruz) and 0.3 nM anti-SDMA (13222, Cell Signaling) antibodies (final concentrations), together with 10 μL of Anti-rabbit IgG (AL104C, Perkin Elmer) and AlphaLISA Acceptor Beads (10 μg/mL final concentration) were added to each well. Plates were then incubated 1 hour at RT. Subsequently 25 μL of the Streptavidin Alpha Donor beads (final concentration: 40 μg /mL) were added to each well in the dark. The final volume within the wells is 50 μL. After incubation for 45 minutes at RT (25°C), plates were read on a Perkin Elmer Envision plate reader.

### Drug Treatments

Methylthioadenosine (MTA), PRMT5 catalytic inhibitor GSK591, and Rapamycin were purchased from Sigma-Aldrich and resuspended in DMSO to 10mM working stocks.

### Protein production

The PRMT5:WDR77 complex was co-expressed in Sf9 cells using the dual promoter pFastBac Dual baculovirus system (Thermo Fisher). Protein purity and molecular mass were confirmed by SDS-Page Coomassie staining and LC/MS. For SPR experiments, untagged full-length PRMT5 was co-expressed with a full-length WDR77 protein containing N-terminal (Strep-Tag II, 3C Protease cleavage site) and C-terminal (ybbR, sortase, His_6_) tags. For crystallography and FP experiments, untagged full-length PRMT5 was co-expressed with a full-length WDR77 protein containing an N-terminal tag (His8, Strep-Tag II, 3C Protease cleavage site) only. The complexes were purified by Streptactin XT affinity resin (IBA). For SPR experiments, the complex was biotin-labeled via incubation with recombinant sortase A5 enzyme (Active Motif) and a custom tri-glycine biotin peptide (Thermo Fisher). The labeled complex was further purified on a Superose 6 size exclusion column (GE Healthcare). For crystallography, the N-terminal tag was removed by incubation with HRV 3C protease (AG Scientific) and further purified by Superose 6 column. PRMT5 proteins were purified and stored in 50 mM HEPES 7.4, 10% glycerol, 1mM TCEP, and 150 mM NaCl. Full-length (aa 2-237) or ΔPBM (aa 2-227) pICln protein was expressed with a N-terminal His6-SUMO tag in BL21(DE3) *E. coli* cells (Thermo Fisher). pICln was first affinity purified by HisTrap column (GE Life Science) in 50 mM Tris pH 8, 500 mM NaCl, 10% glycerol, and 0.5 mM TCEP with imidazole elution. Protein was then purified by gel filtration using a Superdex 200 26/600 GL column (GE Life Sciences) before cleaving the His_6_-SUMO tag with ULP1 enzyme. The tag was removed from solution by passage through a HisTrap column. A monoQ column (GE Life Sciences) was then used as a final purification step to remove degradation products. pICln protein was diluted tenfold into 25 mM Bis-Tris pH 6.0, 0.5 mM TCEP and passed over a monoQ column followed by elution with a linear gradient to 500 mM NaCl. All proteins were quantified by A280 absorbance. pICln protein purity and molecular mass was confirmed by both SDS-Page Coomassie staining and LC/MS.

### Peptides

To generate the fluorescent peptide probe, a synthetic peptide _ac_SRVVPGQFDDADSSDC_am_ was reacted with a rigid maleimide conjugate of KU560 fluorophore (KU Dyes) and purified by reverse phase HPLC. All other peptides were synthesized by Genscript and provided at >95% purity. Net peptide content was determined by nitrogen analysis. Peptides were solubilized in DMSO and stored at −80C.

### Fluorescence polarization

The fluorescence polarization assay buffer (FP buffer) was composed of 10 nM peptide probe, 50 mM HEPES 7.4, 100 mM NaCl, 0.5 mM TCEP, and 0.01% v/v Tween 20. For competition experiments, the PRMT5:WDR77 concentration was fixed at 200 nM protomer. Samples were incubated at room temperature for 30 minutes before reading. Data were collected in black, 384 well non-binding surface plates (Corning) using a Spectramax Paradigm with Rhodamine FP filter set or a Perkin-Elmer Envision with Bodipy TMR FP filter set.

### Crystallography

The PRMT5 complex was crystallized as previously described^31^. To obtain co-crystals, peptide was added to pre-equilibrated drops (without PRMT5 protein) at ~200 μM concentration. This solution was added 1:1 to drops with pre-formed crystals and allowed to incubate for 48 to 72 hr before harvesting directly from drops. Diffraction images were indexed and integrated using XDS^40^. Initial phases for the complex structure were obtained by molecular replacement in Phenix^41^ using PDB 4GQB. Refinement was performed in Phenix and Buster (Global Phasing) with manual building/review in COOT^42^. Surface conservation scores were produced using the Consurf server with automated alignment of all homologs between 50% and 95% sequence identity to the human protein^43^.

### Surface plasmon resonance

Experiments were performed using T200 or S200 Biacore instruments with CM5 sensor chips (GE Healthcare). Sensor chips were immobilized with Neutravidin (Thermo Fisher) at maximum RU via amine coupling. Biotinylated PRMT5 was bound to approximate RU levels of 3000 or 300 for experiments with peptide or full-length pICln as analytes, respectively. For competition experiments, pICln was held constant at a 125 nM concentration and co-injected with varying concentrations of peptide. The assay buffer was 25 mM HEPES pH 7.4, 150 mM NaCl, 0.5 mM TCEP, 0.05% v/v Tween 20, and 1% DMSO. All reported points were measured at binding equilibrium. Data were solvent corrected using a DMSO standard curve and fit with Prism software using a one-site saturation with linear non-specific binding model.

### Peptide enrichment plot

To assess whether the 7-amino acid sequence (herein referred to as “PBM”) was unique to PRMT5 binding proteins, we counted the occurrence of 7-amino acid sequences respectively in PRMT5 interactors and non-interactors (interactors and whole human proteome data was downloaded from BioGrid and Uniprot, respectively) with a sliding window of length 7, also known as 7-mer counting within amino acid context. BLOSUM Clustered Scoring Matrix (BLOSUM100 from R package Biostrings) was taken into consideration to obtain extended 7-mers (one AA difference allowed at a time) when counting the occurrence of a specific 7-mer seed to maximize the flexibility of a potential functional protein pocket^25^. To find co-linear sequences that may form functional protein pockets shared by multiple PRMT5 interactors, more focus was put on 7-mers (as well as its extended 7-mers) found across multiple PRMT5 interactors rather than repetitive sequences from a singular PRMT5 interactor. The fold change (FC) of 7-mer occurrence in interactors with respect to non-interactors was calculated as shown below:

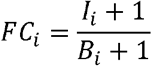

where *I_i_* represents the number of unique interactors containing *i_th_*7-mer seed and its extended 7-mers, *B_i_* represents the number of unique non-interactors containing *i_th_*7-mer seed and its extended 7-mers. A constant of 1 is added to both numerators and denominators to avoid a zero count in the absence of any protein containing a 7-mer seed. Cases where specific 7-mer seeds as well as extended ones are absent in other PRMT5 interactors were first eliminated. Further, considering high sequence similarity found in paralogs, we removed 7-mers found solely in two paralog interactors (e.g. SMARCA4 with SMARCA2 and CDK8 with CDK19). Continuity of fold change was created with the addition of random Gaussian noise for better visualization of the most frequently occurring 7-mer hits.

### SDMA enrichment mass spectrometry

PRMT5 KO-rescue assays were performed as described above. Briefly, 2×10^6 cells stably expressing 3X HA tagged WT, ADA or CD mutants were infected with lentiviral sgRNA targeting PRMT5 in 6 well dishes containing 10 μg/mL polybrene (Santa Cruz). 48 hours after infections, cells were split into 10 cm dishes and selected with puromycin (1 μg/mL) for an additional 48 hours. Following selection, cells were expanded to 3 x 150mm dishes and grown for an additional 5 days. Cells were harvested by scraping and washing 3 times in sterile DPBS. 5% of the cell pellets were set aside to confirm equivalent expression of the PRMT5 constructs and efficient and equivalent knockout of the endogenous PRMT5 across samples. The remaining cell pellets were lysed in PTMScan® Urea Lysis Buffer (20 mM HEPES (pH 8.0), 9.0 M urea, 1 mM sodium orthovanadate (activated), 2.5 mM sodium pyrophosphate, and 1 mM β-glycerol-phosphate). Protein concentration was determined by Bradford assay and 2.8 mg of cell lysate was used from each sample for subsequent analyses.

Mass spectrometry samples were analyzed using the PTMScan method as previously described^44,45^. Briefly, cellular extracts were prepared in urea lysis buffer, sonicated, centrifuged, reduced with DTT, and alkylated with iodoacetamide. 2.8mg total protein for each sample was digested with trypsin and purified over C18 columns for enrichment with the Symmetric Di-Methyl Arginine Motif Antibody (Cell Signaling #13563). Enriched peptides were purified over C18 STAGE tips^46^. Enriched peptides were subjected to secondary digest with trypsin and second STAGE tip prior to LC-MS/MS analysis.

Replicate injections of each sample were run non-sequentially on the instrument. Peptides were eluted using a 120-minute linear gradient of acetonitrile in 0.125% formic acid delivered at 280 nL/min. Tandem mass spectra were collected in a data-dependent manner with a Thermo Orbitrap Fusion™ Lumos™ Tribrid™ mass spectrometer using a top-twenty MS/MS method, a dynamic repeat count of one, and a repeat duration of 30 sec. Real time recalibration of mass error was performed using lock mass (Olsen) with a singly charged polysiloxane ion m/z = 371.101237.

MS/MS spectra were evaluated using SEQUEST and the Core platform from Harvard University^47^. Files were searched against the SwissProt *Homo sapiens* FASTA database. A mass accuracy of +/-5 ppm was used for precursor ions and 0.02 Da for productions. Enzyme specificity was limited to trypsin, with at least one tryptic (K-or R-containing) terminus required per peptide and up to four mis-cleavages allowed. Cysteine carboxamidomethylation was specified as a static modification, oxidation of methionine and mono- or di-methylation on arginine residues were allowed as variable modifications. Reverse decoy databases were included for all searches to estimate false discovery rates, and filtered using a 2.5% FDR in the Linear Discriminant module of Core. Peptides were also manually filtered using a -/+ 5ppm mass error range and presence of a di-methyl arginine residue. All quantitative results were generated using Skyline^48^ to extract the integrated peak area of the corresponding peptide assignments. Accuracy of quantitative data was ensured by manual review in Skyline or in the ion chromatogram files. Mass spectrometry data have been deposited to ProteomeXchange; Dataset identifier PXD019991.

### Mass Spectrometry Data Processing and Analysis

Normalized abundance for 408 peptides containing di-methylated arginines was obtained from mass spectrometry (as described above) in three different PRMT5 variants (WT, ADA and CD), each with two replicates. The missing data in each condition replicate was filled by the minimum values of the replicate MS run. For 3*2 replicates, we performed log2 transformation of the abundance values. Further, the measurements in the WT condition were averaged across two replicates. We zoomed in on 307 peptides with dimethylated “GRG” motif for investigating enriched/depleted substrates. The up- and down-regulated methylation activity of ADA and CD relative to PRMT5^WT^ was quantified by the difference between the average log2 values in WT and log2 transformed values in four replicates. More negative values (blue) in the heatmap represent more enriched methylation activity in the condition replicate. On the other hand, depleted methylation was observed for peptides in each condition replicate when they were annotated with positive values (red). Column hierarchical clustering was performed for four condition replicates while the rows of the heatmap were organized in the order of ascending order of values in ADA replicate 1. Proteins were considered substrates of ADA or CD if they had at least 1 SDMA modified peptide within a “GRG” motif with a FC>2 compared to WT in at least 1 experimental replicate.

**Table 1.**
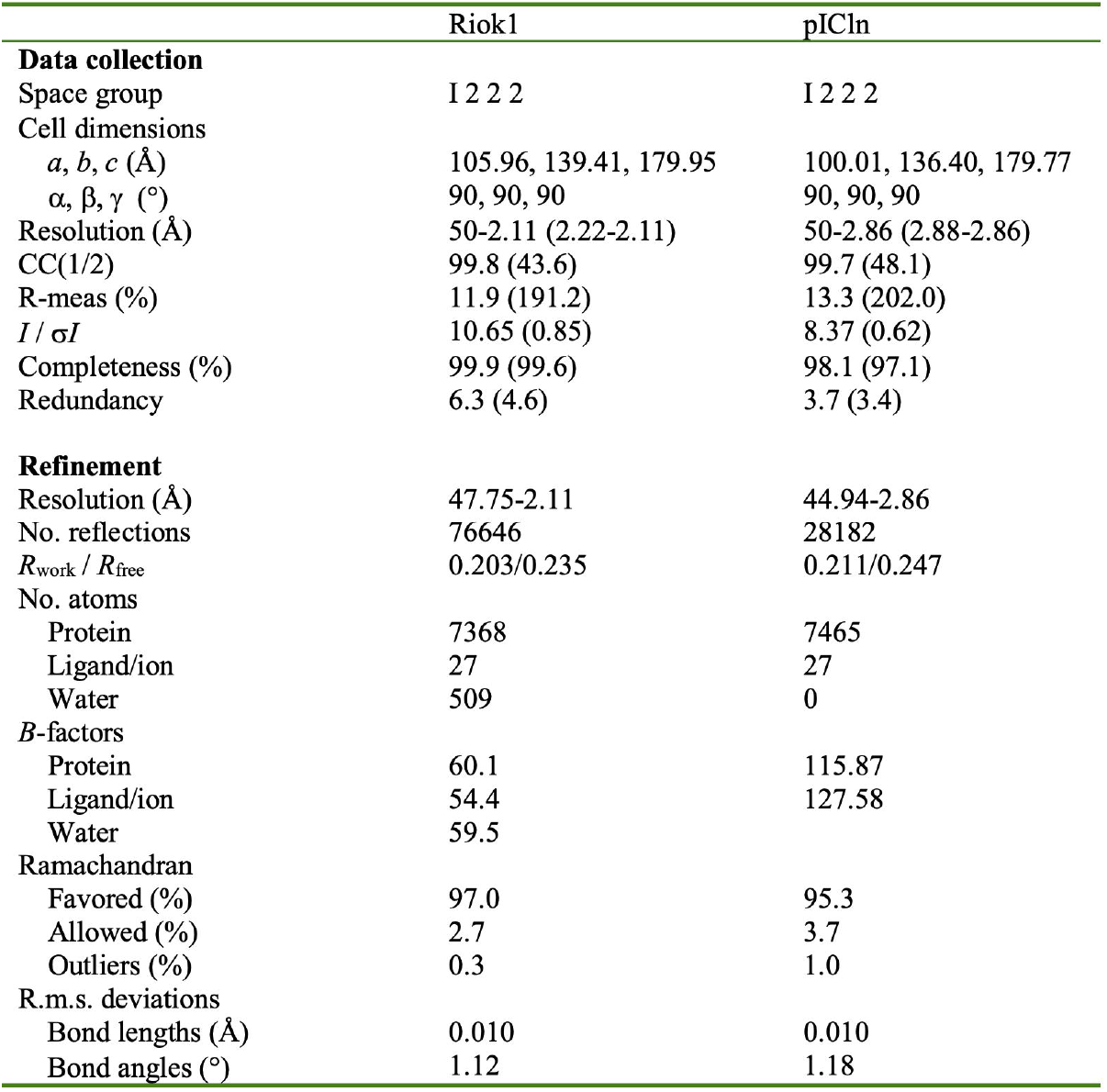
Data collection and refinement statistics.

|  | Riok1 | pICln |
| --- | --- | --- |
| **Data collection** |  |  |
| Space group | I 2 2 2 | I 2 2 2 |
| Cell dimensions |  |  |
| *a*, *b*, *c* (Å) | 105.96, 139.41, 179.95 | 100.01, 136.40, 179.77 |
| $\alpha$, $\beta$, $\gamma$ (°) | 90, 90, 90 | 90, 90, 90 |
| Resolution (Å) | 50-2.11 (2.22-2.11) | 50-2.86 (2.88-2.86) |
| CC(1/2) | 99.8 (43.6) | 99.7 (48.1) |
| R-meas (%) | 11.9 (191.2) | 13.3 (202.0) |
| *I* / $\sigma I$ | 10.65 (0.85) | 8.37 (0.62) |
| Completeness (%) | 99.9 (99.6) | 98.1 (97.1) |
| Redundancy | 6.3 (4.6) | 3.7 (3.4) |
| **Refinement** |  |  |
| Resolution (Å) | 47.75-2.11 | 44.94-2.86 |
| No. reflections | 76646 | 28182 |
| *R*<sub>work</sub> / *R*<sub>free</sub> | 0.203/0.235 | 0.211/0.247 |
| No. atoms |  |  |
| Protein | 7368 | 7465 |
| Ligand/ion | 27 | 27 |
| Water | 509 | 0 |
| *B*-factors |  |  |
| Protein | 60.1 | 115.87 |
| Ligand/ion | 54.4 | 127.58 |
| Water | 59.5 |  |
| Ramachandran |  |  |
| Favored (%) | 97.0 | 95.3 |
| Allowed (%) | 2.7 | 3.7 |
| Outliers (%) | 0.3 | 1.0 |
| R.m.s. deviations |  |  |
| Bond lengths (Å) | 0.010 | 0.010 |
| Bond angles (°) | 1.12 | 1.18 |

## SUPPLEMENTAL DATA

**Supplemental Table 1. X-ray crystal structure data collection and refinement statistics.**

**Supplemental Table 2. PRMT5 methyl-enrichment mass spectrometry results.**

**Supplemental Figure 1.**
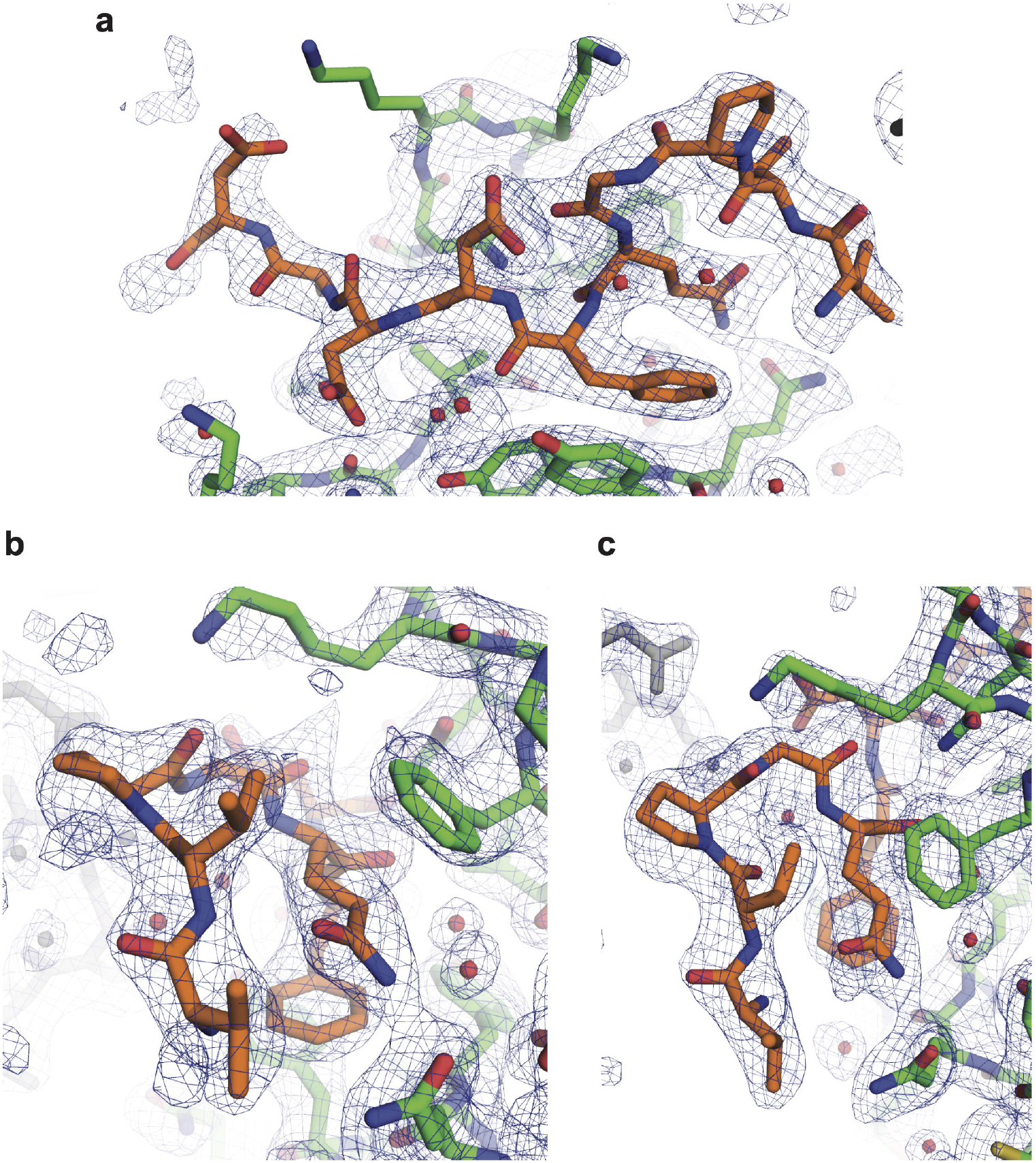
Electron density of the RIOK1-bound X-ray crystal structure. a. a-c. 2Fo-Fc density contoured at 1σ using similar orientation to main Figures 2B-2D

**Supplemental Figure 2.**
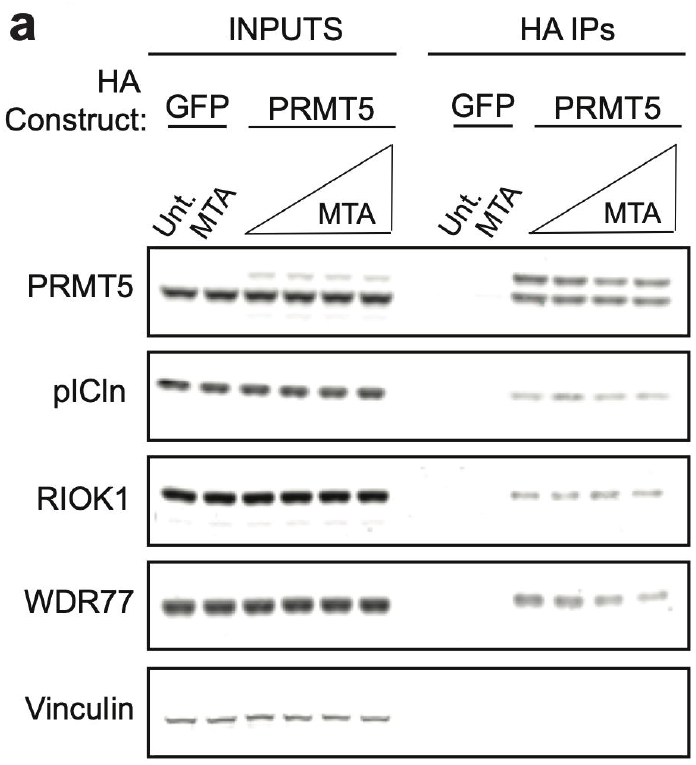
MTA does not affect substrate adaptor recruitment. a. The amount of pICln and RIOK1 bound to PRMT5 in the presence of increasing concentrations of MTA was determined by treating cells for 24h with either vehicle (DMSO), 250 nM, 1 μM, or 10 μM MTA followed by co-IP of HA (PRMT5) and Immunoblotting.

**Supplemental Figure 3.**
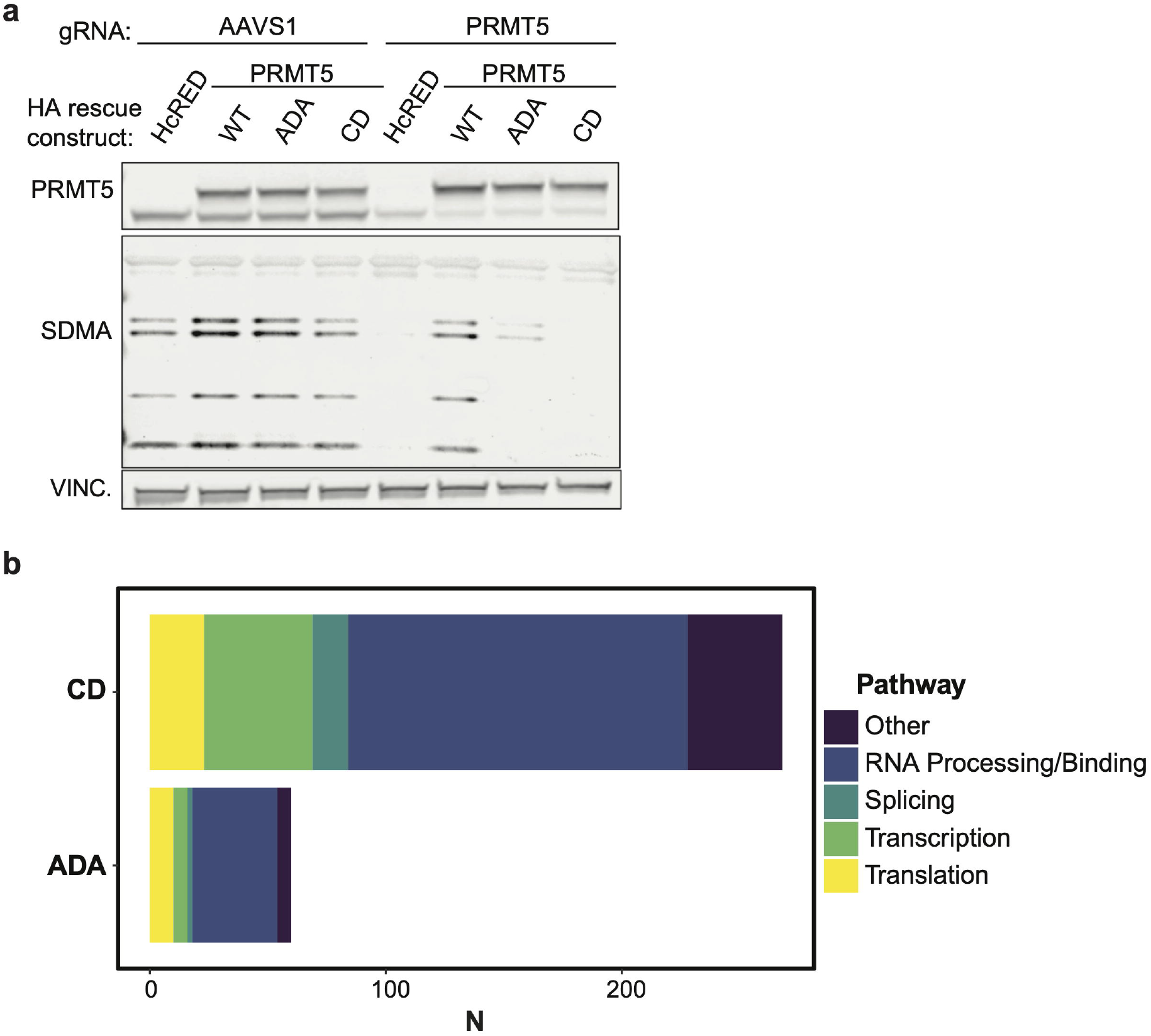
PRMT5^WT^ and PRMT^ADA^ substrates are enriched for RNA binding proteins. a. Immunoblots of a replicate of the lysate samples used to generate Figure 5B-C. Immunoblotting was used to measure the degree of PRMT5-KO and overexpression as well as changes in SDMA levels. Vinculin serves as a loading control. b. Pathway enrichment analysis of substrates changing with CD PRMT5 and ADA PRMT5 compared to WT PRMT5 control rescue (n=number of methyl peptides). Each substrate was assigned to a single GO term.

**Supplemental Figure 4.**
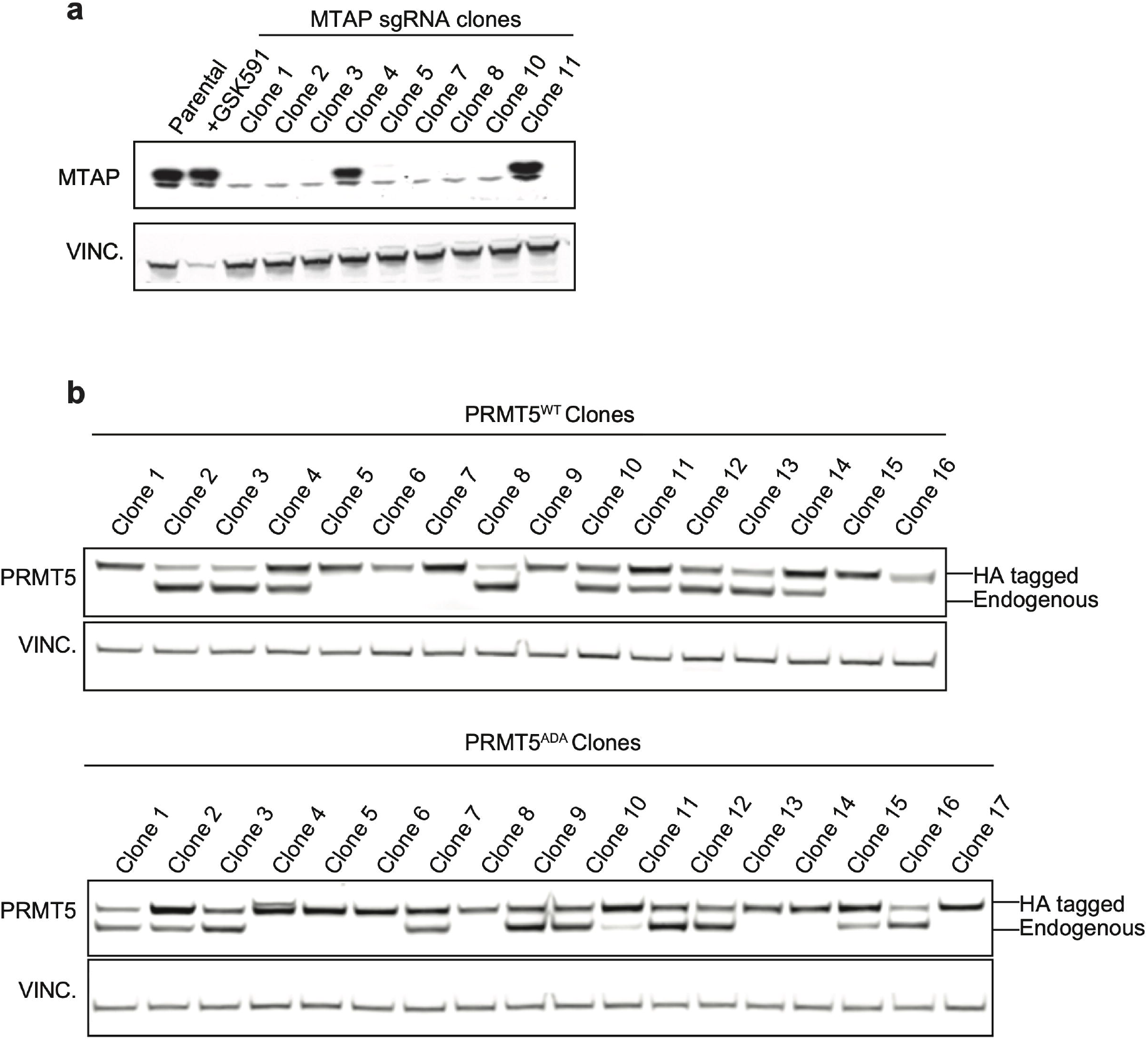
Derivation of MTAP-/- single cell clones and endogenous PRMT5-/- PRMT5^WT^ and PRMTADA cell lines. a. Immunoblotting detected MTAP in each clonal cell line to determine extent of genetic KO following infection of HCT116 with MTAP guide RNA and single cell cloning. Vinculin serves as a loading control. b. Immunoblotting was used to detect endogenous or HA-tagged forms of PRMT5 in each clonal cell line to determine extent of genetic KO of endogenous PRMT5 following infection with PRMT5 guide RNA in the HCT116 MTAP KO pool (comprised of a pool of clones from (A)) which were also stably expressing PRMT5^WT^ or PRMT5^ADA^. Vinculin serves as a loading control.

## Acknowledgements

We thank the following core facilities and platforms at the Broad Institute for support of our work: Genetic Perturbation Platform, Flow Cytometry Core, Proteomics Platform, Center for the Development of Therapeutics (CDoT), and Cancer Program. We thank Kathryn Drezelo for operational support. This work was supported by an NIH F32 (CA232543-01) to KMM and NIH R01 (1R01CA233626-01A1) to WRS. This work was also supported by the Deerfield-Broad Discovery Research Collaboration.

## Contributions

K.M., W.R.S., B.M., D.P., and A.I. designed all of the experiments. K.M., B.M., C.B., and N.A. performed all of the experiments. V.Y. and M.S. performed the mass spectrometry analysis. R.L. and S.J. performed the peptide enrichment analysis. R.L. also performed the statistical analysis for the mass spectrometry dataset. Y.F., F.K.B., and A.S. performed or provided experimental support for PRMT5 AlphaLISA and NanoBiT assays. M.O., M.R., and Y.B. provided experimental support for PRMT5 *in vitro* experiments. D.R. assisted in refinement of the crystal structure. K.M., B.M., and W.R.S. wrote the manuscript. All authors have discussed the results and provided comments on the manuscript.

## Notes

http://www.proteomexchange.org/

https://www.wwpdb.org/pdb?id=pdb_00006v0o

### Competing Interest Statement

Materials related to this article have been included in a provisional patent application by the Broad Institute. W.R.S. is a Board or SAB member and holds equity in Peloton Therapeutics, Ideaya Biosciences, Civetta Therapeutics, and Bluebird and has consulted for Array, Astex, Dynamo Therapeutics, Ipsen, PearlRiver Therapeutics, Sanofi, and Servier; and receives research funding from Pfizer Pharmaceuticals and Deerfield Management.

